# Exploration of the structural and functional diversity in the metamorphic RfaH subfamily

**DOI:** 10.64898/2026.03.16.712203

**Authors:** Cyndi Tabilo-Agurto, Bastián González-Bustos, Javiera Reyes, Bing Wang, Damaris Palomera, Verónica Del Río-Pinilla, Camila Neira-Mahuzier, Valentina Vera-Sandoval, Irina Artsimovitch, Pablo Galaz-Davison, César A. Ramírez-Sarmiento

## Abstract

RfaH, a member of the universally conserved NusG/Spt5 family, activates transcription and translation of virulence genes by bridging the transcribing RNA polymerase to the translating ribosome. Unlike its monomorphic, constitutively active paralog NusG, RfaH fold-switches from an autoinhibited state, wherein its C-terminal domain forms an α-helical hairpin bound to the N-terminal domain, into an active NusG-like state through C-terminal domain dissociation and refolding into a β-barrel. RfaH has been proposed to evolve from NusG via gene duplication by first activating genes adjacent to its production site, before developing autoinhibition to selectively control distant genes. Using AlphaFold2, we predicted the structures of thousands of RfaH homologs and identified a subset (∼14%) that appear predominantly folded in the active state. *In vivo* assays confirmed that these putative monomorphic homologs exhibit constitutive activity comparable to known RfaH mutants. Phylogenetic and genomic analyses revealed that these monomorphic proteins form a distinct clade and are preferentially located within or next to long virulence-associated operons. Together, these results further support a stepwise evolutionary model of RfaH specialization through structural transformation.

## INTRODUCTION

Metamorphic proteins are characterized by their reversible fold-switch between two disparate, yet stable native states encoded by a single amino acid sequence, necessary for regulating or encoding different biological functions (Porter *et al*, 2024). These proteins have been discovered by solving their three-dimensional structures under different contexts, such as sequence truncations (Bullough *et al*, 1994; Burmann *et al*, 2012), residue substitutions that stabilize their different folds (Chang *et al*, 2015; Cai *et al*, 2020) or changes in environmental conditions (Littler *et al*, 2004; López-Pelegrín *et al*, 2014), with rare cases in which both native states are observed in a single experimental structure (Mapelli *et al*, 2007).

Over 100 proteins across all kingdoms of life have been described to switch folds (Porter & Looger, 2018), with recent discoveries in viral proteins suggesting that these numbers will increase (Cai *et al*, 2020; Arnold *et al*, 2026). In fact, bioinformatic analysis of folding cooperativity and ambiguous secondary structure predictions from sequence information suggest that ∼4% of the proteins available in the PDB may switch folds (Porter & Looger, 2018).

While computational and experimental biophysics have shed light on the fold-switching mechanisms of several metamorphic proteins, such as *Escherichia coli* RfaH (Zuber *et al*, 2019; González-Higueras *et al*, 2024; Tabilo-Agurto *et al*, 2025; Cai *et al*, 2025), cyanobacterial KaiB (Chang *et al*, 2015; Rivera *et al*, 2022), and human XCL1 (Tuinstra *et al*, 2008; Camilloni & Sutto, 2009; Tyler *et al*, 2011), less is known about the evolutionary emergence of these proteins in nature. To date, only the emergence of XCL1 has been thoroughly explored by ancestral sequence reconstruction and structural characterization by nuclear magnetic resonance spectroscopy of these ancestors, suggesting that this protein evolved from an ancestor with a monomeric chemokine fold into a novel dimeric native state (Dishman *et al*, 2021). Overall, the consensus is that most, if not all, metamorphic proteins evolved by encoding an alternative native state within monomorphic protein families with a canonical fold (Dishman & Volkman, 2023; Porter *et al*, 2024).

The quintessential metamorphic virulence factor RfaH (Artsimovitch & Ramírez-Sarmiento, 2022) is another well-studied protein in terms of its evolution, with a concrete hypothesis of its emergence within the universally conserved NusG/Spt5 family of transcription elongation factors (Tomar & Artsimovitch, 2013). RfaH arose through gene duplication of NusG (Bailey *et al*, 1997), a protein that in *E. coli* is associated with RNA polymerase (RNAP) transcribing almost all genes (Mooney *et al*, 2009a). NusG is a monomorphic protein composed of two domains, the NusG N-terminal domain (NTD), which binds to RNAP in transcription elongation complexes (TEC), and the C-terminal Kyrpides, Ouzounis, Woese domain (CTD) folded as a β-barrel (βCTD) that binds to either Rho or the ribosome (Mooney *et al*, 2009b; Burmann *et al*, 2010). NusG enables RNAP to monitor the RNA quality and cellular conditions by either promoting early transcription termination of foreign, antisense, or damaged RNAs by Rho (Cardinale *et al*, 2008; Peters *et al*, 2012), or by supporting transcription-translation coupling (Wang *et al*, 2020b; Webster *et al*, 2020).

In contrast, the *E. coli* paralog RfaH, with only 21% sequence identity to NusG, is a metamorphic protein. In its resting, autoinhibited state, the RfaH-CTD is folded as an α-helical hairpin (αCTD) tightly bound to the RNAP-binding site on the NTD (Burmann *et al*, 2012; Zuber *et al*, 2018). Despite a 100-fold cellular excess of NusG (Schmidt *et al*, 2016), RfaH excludes NusG from RNAP (Belogurov *et al*, 2009) due to its 10-fold higher affinity (Kang *et al*, 2018). In *Enterobacteriaceae*, RfaH dramatically activates expression of several long horizontally-acquired operons related to bacterial virulence (capsule, cell wall, toxins, adhesins, pilus and lipopolysaccharide [LPS] biosynthesis); these operons frequently lack ribosomal binding sites (RBS) and are silenced by the joint action of Rho and nucleoid-associated proteins (Hustmyer *et al*, 2022; Wang *et al*, 2022). RfaH is recruited to RNAP at a specific 12-nt sequence known as *operon polarity suppressor* (*ops*) (Bailey *et al*, 1997) that forms a short hairpin in the non-template DNA strand on the surface of a paused TEC (Artsimovitch & Landick, 2002; Zuber *et al*, 2018). The RfaH interactions with *ops* and RNAP triggers activation of RfaH by interdomain dissociation and fold-switching of its αCTD into the canonical βCTD of NusG (Burmann *et al*, 2012; Zuber *et al*, 2024). Once RfaH unbinds from the TEC, it folds back into its autoinhibited state (Zuber *et al*, 2019). While RfaH-CTD does not bind Rho (Lawson *et al*, 2018), it interacts with the ribosomal protein S10 to recruit the ribosome to RBS-less mRNAs (Burmann *et al*, 2012) and may subsequently couple transcription and translation (Molodtsov *et al*, 2024). By simultaneously excluding NusG and anchoring the ribosome, RfaH ensures the expression of distal genes within long *ops*-containing virulence operons.

It has been hypothesized that after *nusG* gene duplication, monomorphic ancestral RfaH with the canonical NusG fold (βRfaH) first lost its binding to Rho and solubility, while gaining higher affinity for RNAP to regulate genes closely linked to its production site (i.e., in *cis*), impeding displacement by the more abundant NusG protein (Belogurov *et al*, 2009; Wang *et al*, 2020a). Then, RfaH would have evolved to gain DNA sequence specificity and solubility by folding into the autoinhibited state (αRfaH) to enable its action in *trans* over operons far from the genomic loci encoding RfaH (Belogurov *et al*, 2009; Wang *et al*, 2020a). However, there is neither direct or indirect evidence of extant, functional monomorphic βRfaH proteins, nor of their in-*cis* regulation.

Deep learning (DL) protein structure prediction methods such as AlphaFold2 (AF2) only predict one of the two native states of fold-switching proteins (Chakravarty & Porter, 2022; Chakravarty *et al*, 2024) However, different results are obtained when reducing the number of homologous protein sequences in the multiple sequence alignment (MSA) generated as input during the process (Wayment-Steele *et al*, 2024; Guan *et al*, 2024; Lee *et al*, 2025). Nevertheless, AF2 in its default settings predicts the autoinhibited state for *E. coli* RfaH and for the distant homolog from *Vibrio cholerae*, which only shares 43% sequence identity (Artsimovitch & Ramírez-Sarmiento, 2022).

Based on the assumption that predicting the autoinhibited state is indicative of their metamorphic behavior, one can hypothesize that AF2 could be used to predict fold-switching RfaH orthologs. Conversely, predicting the canonical NusG-like state, which has been achieved using evolutionary-based and physics-based mutations of interdomain CTD residues in *E. coli* RfaH – some of which have been experimentally validated to be constitutively active in previous studies (Shi *et al*, 2017; Freiberger *et al*, 2023; Tabilo-Agurto *et al*, 2025) – would suggest potential monomorphic candidates. This opens the door to explore the evolutionary hypothesis of the structural and functional emergence of the RfaH fold-switch within the NusG/Spt5 family.

Based on these precedents, and also on recent discussions that fold-switching proteins are proof of protein folds existing in a continuum of states (Tian & Best, 2020; Porter, 2023; Kulkarni *et al*, 2025), we used ColabFold (Mirdita *et al*, 2022), a cloud-based implementation of AF2 (Jumper *et al*, 2021), to perform a large-scale structure prediction of *E. coli* RfaH homologs with a range of sequence identity from several sequence databases. While ∼66% of the ∼20,000 RfaH structures were predicted to have an αCTD, ∼14% of the sequences of RfaH homologs (22% of the predicted structures) were mainly predicted in the β-fold, suggesting they constitute potential monomorphic RfaH proteins, and 9% were predicted to have mixed α/β CTDs. These structures were observed in targeted molecular dynamics (TMD) (Schlitter *et al*, 1994) simulations of the *E. coli* RfaH fold-switch and were assessed as reliable by the AF2-based structure scoring framework AF2Rank (Roney & Ovchinnikov, 2022), suggesting that they may represent evolutionary and structural intermediates of RfaH fold-switching (Figure 1).

**Figure 1.**
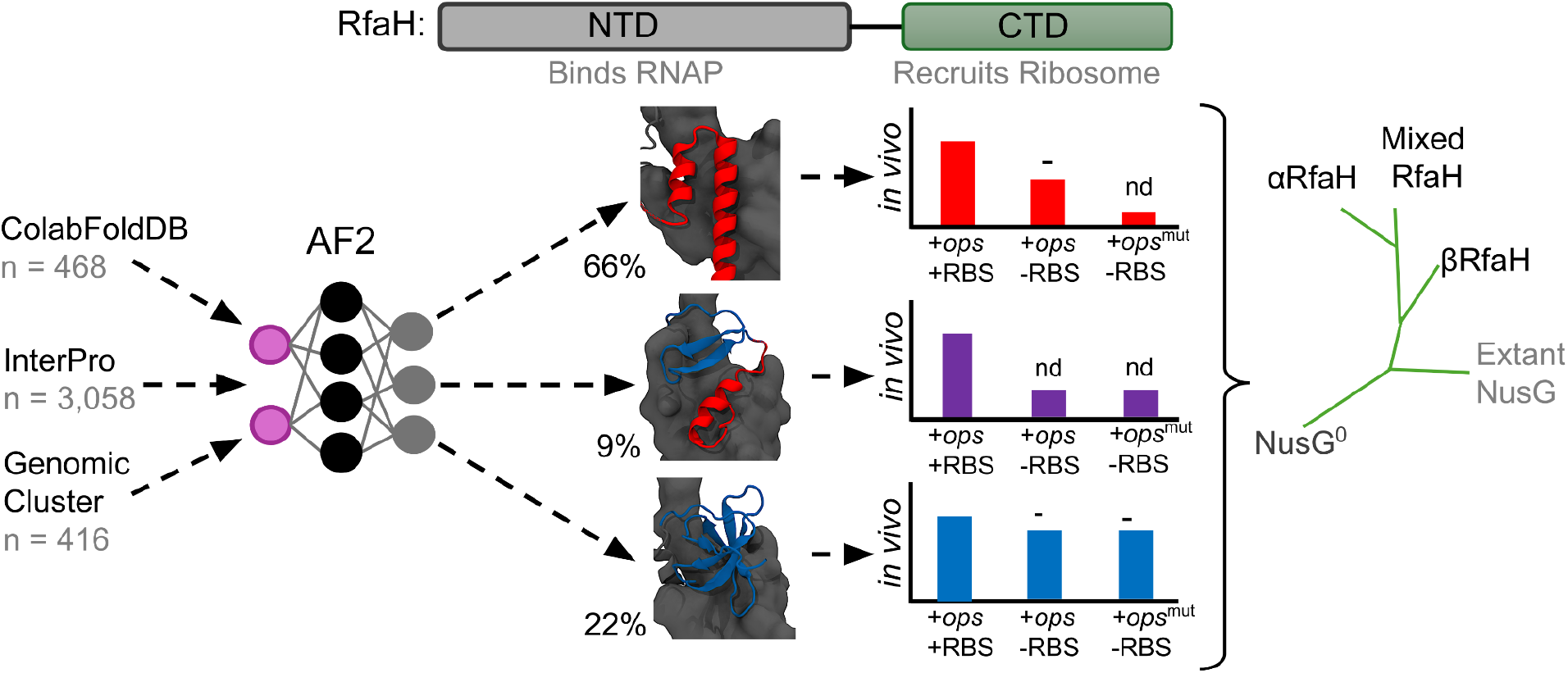
AF2-, activity-, and phylogeny-guided classification of the structural and functional diversity of extant RfaH homologs. (**Top**) Architecture of RfaH displaying the NTD and the fold-switching CTD connected by a linker. (**Bottom**) Scheme of the computational and experimental approach to explore the structural and functional diversity of extant RfaH proteins. The structures of >3,900 sequences from different databases were predicted using AF2 in ColabFold, and classified based on the secondary structure content of their CTD into either αCTD, mixed α/β CTD, or βCTD folds. Three representative RfaH homologs from the Genomic Cluster predicted in each state were subjected to RfaH-dependent *in vivo* transcription-translation assays to explore the effects of the *ops* signal and the RBS on their activity using a luminescence reporter. Lastly, a maximum likelihood phylogenetic analysis grouped these different extant proteins in separate clades for metamorphic αRfaH, monomorphic βRfaH, and RfaH with mixed α/β CTD, matching their structural and functional diversity. Nd = not detectable (similar to luminescence with condition lacking RfaH expression).

We selected nine RfaH homologs from a database of RfaH sequences with known genomic origin, whose predicted structures exhibited either αCTD, βCTD or mixed α/β CTD, for experimental testing using *in vivo* translation assays. Proteins with mixed α/β CTD fold did not display RfaH-dependent transcription-translation coupling, whereas monomorphic βRfaH proteins behaved similarly to the constitutively active E48A RfaH variant that does not require the *ops* for its activity (Burmann *et al*, 2012). Lastly, phylogenetic inference and genomic analysis of RfaH homologs and their predicted structures revealed that monomorphic βRfaH proteins cluster into a defined clade and are located next to pathogenicity-related long operons in their corresponding genomes, in contrast to metamorphic αRfaH homologs which operate in *trans*. Together, these results provide further evidence that RfaH evolved from monomorphic, NusG-like ancestors that acted locally near their production sites, a lineage that appears to remain extant in some species (Figure 1).

## RESULTS

### AF2 predicts multiple conformations for different RfaH homologs

Amino acid sequences for *E. coli* RfaH homologs were retrieved from ColabFold database (ColabFoldDB) (Mirdita *et al*, 2022), corresponding to 468 sequences after sequence filtering (see *Methods*); InterPro (Blum *et al*, 2025) (family entry IPR010215) with 3,058 sequences, and a previously reported list of 416 RfaH sequences (Cluster 1) curated by hidden Markov models (HMM) with known genomic context that were classified by Markov clustering and include *E. coli* RfaH (Genomic Cluster, hereafter) (Wang *et al*, 2020a).

These 3,942 sequences were used as input for DL-guided protein structure prediction using AF2 with the default ColabFold protocol, their TM-score (Zhang & Skolnick, 2004) (a normalized structure superimposition quality metric) and the RMSD against the αCTD and βCTD states from *E. coli* RfaH was assigned using TM-align (Zhang & Skolnick, 2005) and the secondary structure of their CTDs with STRIDE (Frishman & Argos, 1995) (see *Methods*). We employed this analysis to determine secondary structure thresholds that enabled us to distinguish between metamorphic RfaH (αRfaH), monomorphic RfaH (βRfaH) or RfaH with mixed α/β secondary structure in the CTD (Appendix Figure S1). Since proteolytic removal of the NTD (Burmann *et al*, 2012) and ColabFold predictions (Artsimovitch & Ramírez-Sarmiento, 2022) have shown that the isolated CTD assumes the β fold, predictions were also made using the isolated CTDs of the proteins from each database to test whether potential metamorphic proteins indeed switch folds (Appendix Figure S2).

Analysis of the 19,710 generated structures showed that AF2 predicted the autoinhibited state of RfaH in sequences from all three databases, although with different propensities (Figure 2 and Appendix Figure S2). For the predicted structures from ColabFoldDB, which includes some representatives that are more distant to *E. coli* RfaH (mean sequence identity to *E. coli* RfaH = 31.5%), 33% of all 5 structures generated per sequence were predicted to fold as αRfaH, whereas 40% of the predicted structures had a βCTD fold resembling that of NusG (Figure 2A and Appendix Figure S2A and S2C). These βRfaH structure predictions strongly suggest that these homologs might be monomorphic. Also, ∼14% of the ColabFoldDB predicted structures of full-length RfaH homologs exhibited a CTD with mixed α/β secondary structure (Figure 2A and 2E and Appendix Figure S2E). It is worth noting that about 12% of the predicted structures from ColabFoldDB were uncategorized due to their lack of secondary structure.

**Figure 2.**
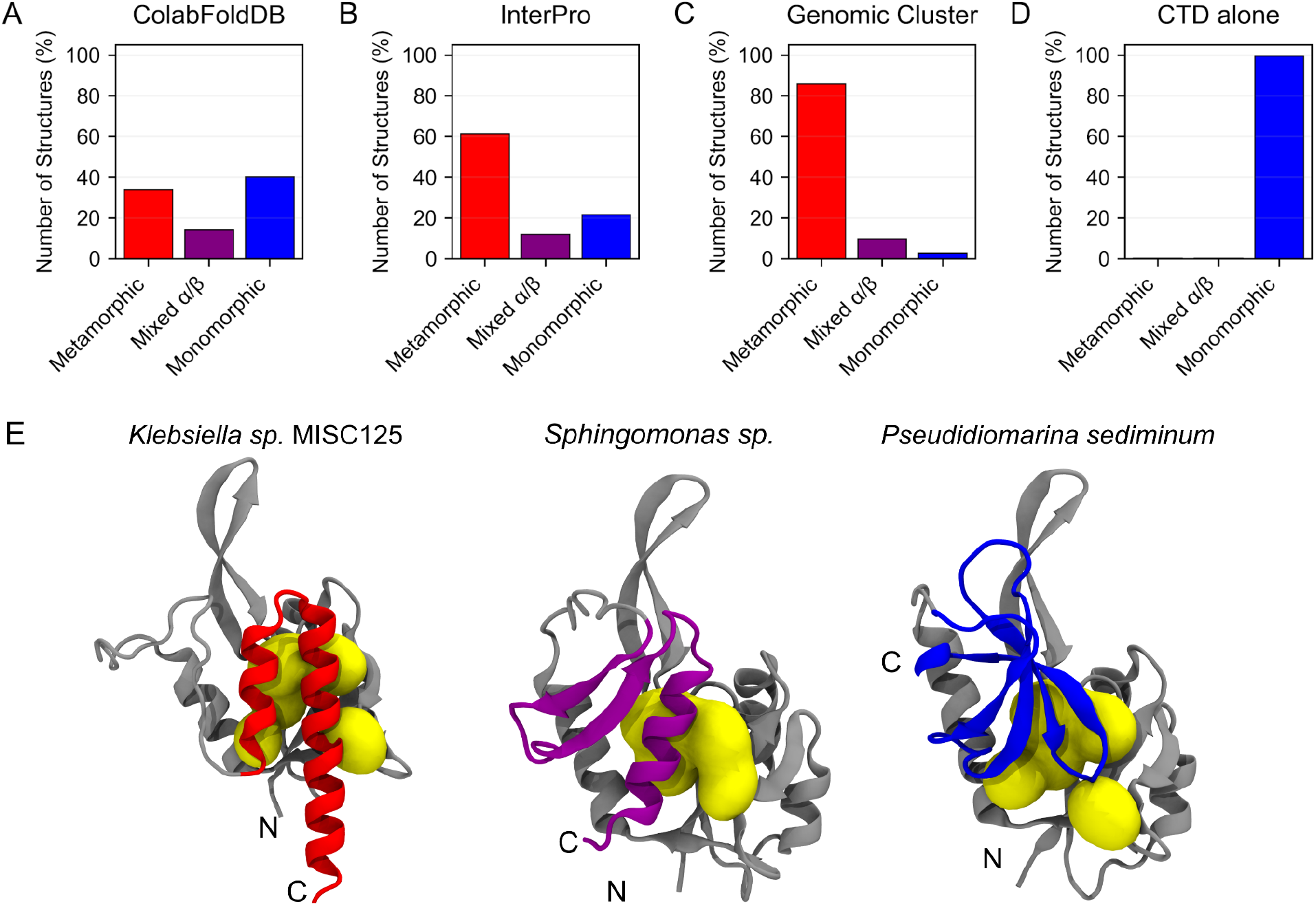
Distribution of predicted structures of extant RfaH homologs with αCTD, βCTD or mixed secondary structure from the ColabFoldDB (**A**), InterPro (**B**) and Genomic Cluster (**C**) databases. The secondary structure associated with the CTD region was ascertained using STRIDE and the CTD region was selected based on a sequence alignment to *E. coli* RfaH as reference. About 66% of the predicted structures of full-length RfaH homologs from ColabFoldDB (**A**) and InterPro (**B**) exhibit an αCTD, while ∼22% are predicted with a βCTD and ∼9% are predicted to have mixed α/β secondary structure. Most of the RfaH sequences from the Genomic Cluster (**C**) are predicted to fold into the αCTD, and when the isolated CTD from these sequences is used as input for protein structure prediction (**D**), they fold into the canonical βCTD of the NusG family. (**E**) Cartoon representations of representative RfaH homologs from the Genomic Cluster database exhibiting an αCTD (red), βCTD (blue) or mixed secondary structure (purple). The structurally conserved NTD is shown in gray, and interdomain contacts are shown in yellow.

The fraction of predicted structures with mixed α/β secondary structure is similar to ColabFoldDB in the case of the InterPro database, with 12% of the predicted structures exhibiting these patterns (Figure 2B and Appendix Figure S2B), suggesting that the prediction of a mixed α/β CTD for some RfaH homologs is not rare for AF2. The predicted structures of the InterPro sequences also show an important increase from 34% structures predicted from ColabFoldDB in the αRfaH fold to 61% in InterPro, along with a decrease in the number of structures with a βRfaH fold from 40% to 22%. This is likely due to filtering out distant NusG-like proteins in the InterPro database (mean sequence identity to *E. coli* RfaH = 49.4%) that might be still present in ColabFoldDB, similar to previous articles where NusG-like sequences were filtered out based on sequence-based secondary structure predictions before coevolutionary analysis (Galaz-Davison *et al*, 2022).

Finally, the predicted structures for the sequences from the Genomic Cluster exhibited the highest fraction of structures with an αRfaH fold (86%), with only 10% having a mixed α/β secondary structure content in their CTD, and merely 3% exhibiting a βRfaH fold (Figure 2C and Appendix Figure S2C). This is a strong indicator that this set is indeed curated towards close homologs of *E. coli* RfaH, even though a small percentage of these orthologs are predicted by AF2 as monomorphic, most likely due to the higher sequence identity of these orthologs with *E. coli* RfaH (mean sequence identity of 53.0%).

To determine if the predicted structures with an αRfaH fold from the different databases correspond to metamorphic representatives of this NusG subfamily, we performed two analyses. First, the structure of the isolated CTD of each protein was also predicted using ColabFold. With a handful exceptions, most of the isolated CTD sequences were predicted by AF2 to fold into the canonical βCTD structure (Figure 2D and Appendix Figure S2), with 99% of them containing only β-strands.

Second, we determined the Cβ-Cβ sidechain distance in all predicted structures of full-length RfaH between residues equivalent to the interdomain E48-R138 salt bridge present in *E. coli* RfaH (see *Methods*). This salt bridge is known to stabilize the autoinhibited state, and its disruption via the E48A mutation leads to a 1:1 equilibrium between the autoinhibited and active state (Burmann *et al*, 2012). This analysis confirmed that the median and 25% lower interquartile distances for the predicted structures with an αRfaH fold from all databases were similar or slightly above the Cβ-Cβ sidechain distance between E48 and R138 (9.56 Å) in the best ranked predicted structure of *E. coli* αRfaH using ColabFold (Figure 3).

**Figure 3.**
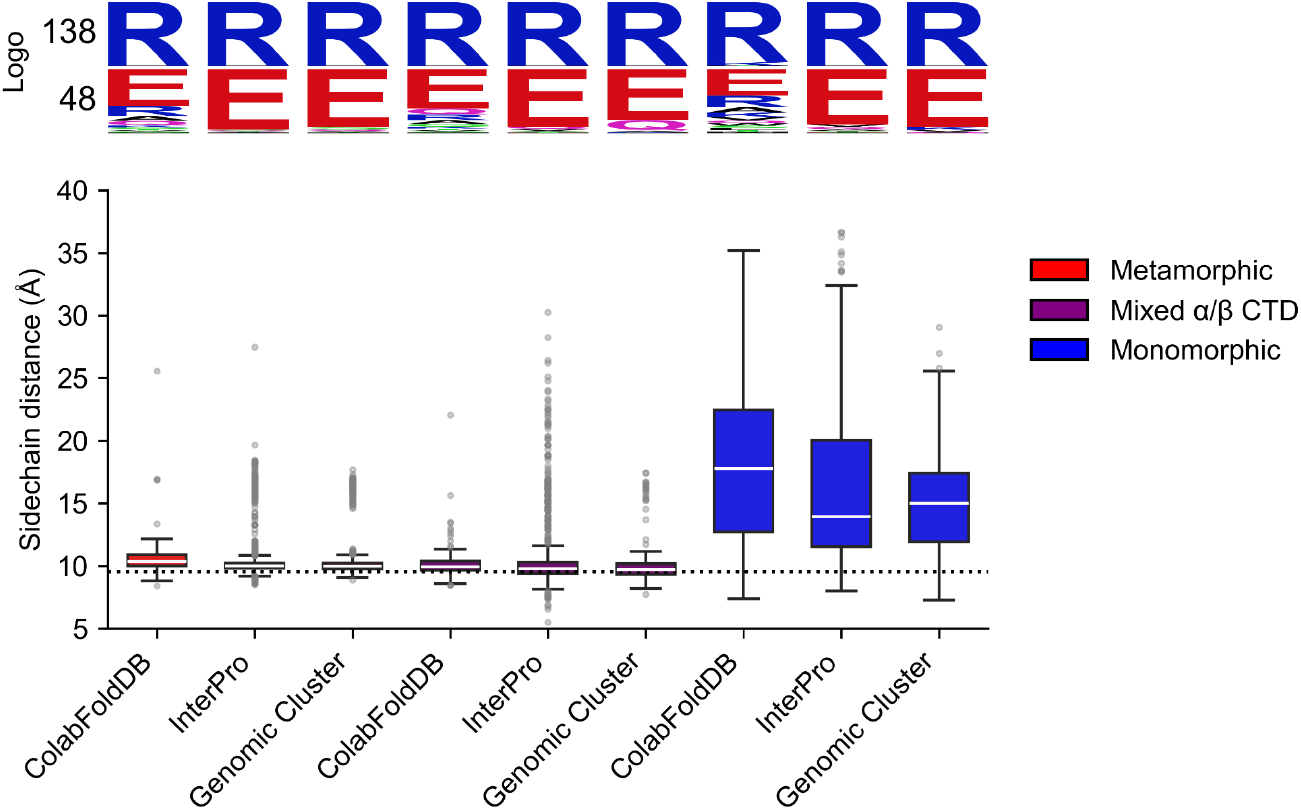
Cβ-Cβ sidechain distance between residues equivalent to the *E. coli* RfaH E48-R138 interdomain salt bridge in all predicted structures. For every structure prediction in each sequence set, residues in equivalent positions to E48 and R138 in *E. coli* RfaH, which are involved in a salt bridge that is critical for the stability of its autoinhibited state, were identified by pairwise sequence alignment to *E. coli* RfaH. Then, the Cβ-Cβ distance between these selected residues in each structure was determined. The box plots show the side chain distance distribution for each AF2-predicted dataset split by their classification (metamorphic in red, first 3 box plots on the left side; mixed α/β in purple, 3 box plots in the middle; monomorphic in blue, last 3 box plots on the right side). The sequence conservation logo of the residues involved in these interactions is displayed on top. The median is shown as a horizontal white line in each box plot, the lower and upper ends indicate the 25% and 75% interquartile range, and the whiskers extend to the percentile 1 and 99, with points representing outliers. The dotted line represents the Cβ-Cβ distance for the E48-R138 salt bridge in the best ranked AF2 structure of *E. coli α*RfaH. The number of analyzed structures to generate this plot is 19,555.

For the predicted structures in the αRfaH fold from ColabFoldDB, InterPro and Genomic Cluster, the median distances correspond to 10.34, 10.01 and 9.97 Å, respectively, and the distances in the 25% lower interquartile range are 10.00, 9.80 and 9.75 Å, respectively. For the structures predicted with a mixed α/β CTD from ColabFoldDB, InterPro and Genomic Cluster, the median distances were 9.93, 9.78 and 9.70 Å, respectively, and the distances in the 25% lower interquartile range are 9.64, 9.41 and 9.32 Å, respectively, which are even shorter than the best ranked predicted structure of *E. coli* αRfaH using ColabFold.

In contrast, the AF2-predicted structures of full-length RfaH homologs with a βCTD fold have median distances of 17.76 Å for ColabFoldDB, 13.96 Å for InterPro and 15.00 Å for Genomic Cluster, and the distances in the 25% lower interquartile range are 12.71, 11.55 and 11.91 Å, respectively. Notably, these differences are observed regardless of the high conservation of the glutamic acid and arginine residues across all sequences in positions equivalent to *E. coli* RfaH residues 48 and 138, respectively (Figure 3).

Importantly, these results are well aligned with the differences in sequence identity for the predicted structures of RfaH homologs in their different structural classifications (Appendix Figure S3). Upon retrieving the best ranked predicted protein structures using the average per-residue local distance difference test (pLDDT, a confidence metric from AF2) (Jumper *et al*, 2021) of their CTDs as a proxy, we observed that the predictions of the αCTD and the βCTD folds exhibit higher pLDDT than the predictions of the CTD structures with mixed α/β secondary structure (Appendix Figure S4).

Specifically, the median pLDDT for the αRfaH and βRfaH structures are 63.12 and 68.28 for ColabFoldDB, 62.29 and 62.39 for InterPro, and 61.16 and 56.16 for Genomic Cluster, respectively. Moreover, the upper 75% interquartile range has values of 67.26 and 80.85 for ColabFoldDB, 63.56 and 77.39 for InterPro, and 62.97 and 60.47 for Genomic Cluster, respectively. In contrast, the predictions of the mixed α/β have lower median and 75% interquartile range values of 48.84 and 56.66 for ColabFoldDB, 47.21 and 52.79 for InterPro, and 44.61 and 48.85 for Genomic Cluster, respectively (Appendix Figure S4).

When analyzing the sequence identity percentage per structural classification against *E. coli* RfaH, the best αRfaH predicted structures from the ColabFoldDB, InterPro and Genomic Cluster databases have a median of 31.5, 47.5 and 48.7% sequence identity to *E. coli* RfaH, respectively. Conversely, the βRfaH predicted structures have a median of 29.6% for ColabFoldDB, 34.6% for InterPro and 38.9% for Genomic Cluster. Moreover, the 75% upper interquartile range has higher value for the αRfaH predicted structures from InterPro and Genomic Cluster than those from the βRfaH predicted structures (Appendix Figure S3).

Altogether, these results strongly suggest that the predicted structures from all databases in the αRfaH fold represent the autoinhibited state unique to metamorphic RfaH homologs, whereas those predicted in the βRfaH fold represent putative extant monomorphic RfaH homologs.

### A mixed α/β CTD topology is observed in fold-switching simulations of *E. coli* RfaH

An interesting finding from the large-scale protein structure prediction of RfaH homologs from ColabFoldDB, InterPro and the Genomic Cluster is the significant number of predicted structures with mixed α/β CTD (Figure 2 and Appendix Figure S2) in close proximity to the NTD, as ascertained by the sidechain distance between residues equivalent to the E48-R138 salt bridge in *E. coli* αRfaH (Figure 3). Moreover, predictions of the isolated CTD for RfaH homologs from all three databases mostly comprise β-strands (Appendix Figure S2). These results, and our previous work using ColabFold to predict the structures of alanine scanning mutants of interdomain CTD residues in full-length *E. coli* RfaH, which also identified structures with such mixed α/β content (Tabilo-Agurto *et al*, 2025), strongly suggest that the mixed α/β topology originates from coevolutionary interactions retrieved from the MSA between NTD and CTD residues, rather than from coevolutionary signals within the CTD alone.

We sought to determine whether the mixed α/β CTD structures predicted by AF2 are also observed during the fold-switch of *E. coli* RfaH. To pursue this, we performed TMD (Schlitter *et al*, 1994), an out-of-equilibrium simulation technique, to explore rare conformational transitions, such as the fold-switch of *E. coli* RfaH CTD in the context of the full-length protein, using an external potential over a collective variable that accurately describes this change in structure (see *Methods*). Then, we extracted all the structural ensembles explored during these TMD trajectories for analysis using AF2Rank (Roney & Ovchinnikov, 2022), a framework that repurposes AF2 as a scoring function to evaluate and rank the quality of protein structures.

For this experiment, ensembles explored by 600 TMD trajectories of 100 ps from αRfaH to βRfaH and vice versa (300 in each direction) using implicit solvent (Appendix Figure S5) were first clustered based on Principal Component Analysis (Appendix Figure S6) to select the best 100 representative trajectories of the *E. coli* RfaH fold-switch in each direction (see *Methods*), and analyzed alongside the minimized structures of the *E. coli* αRfaH and βRfaH using AF2Rank after cycling once through the EvoFormer and Structure Module of AF2. These simulations were so short and were made using a range of forces so broad that only a small fraction of them reached the target state and are not amenable for thermodynamic analysis. However, their explored ensembles constitute a set of ∼7,500 decoy structures from fold-switching simulations for calculating confidence metrics of the input structures, such as the predicted Template Modelling score (pTM) and mean pLDDT (Jumper *et al*, 2021), while also obtaining the minimized output structure after AF2 refinement based on its learned potential energy function (Roney & Ovchinnikov, 2022). This approach enables us to address how much these decoys resemble the mixed α/β CTD structures predicted by AF2 from our sequence databases and whether the learned energy function from AF2 converges to a subset of output structures from this set of initial decoys (Figure 4).

**Figure 4.**
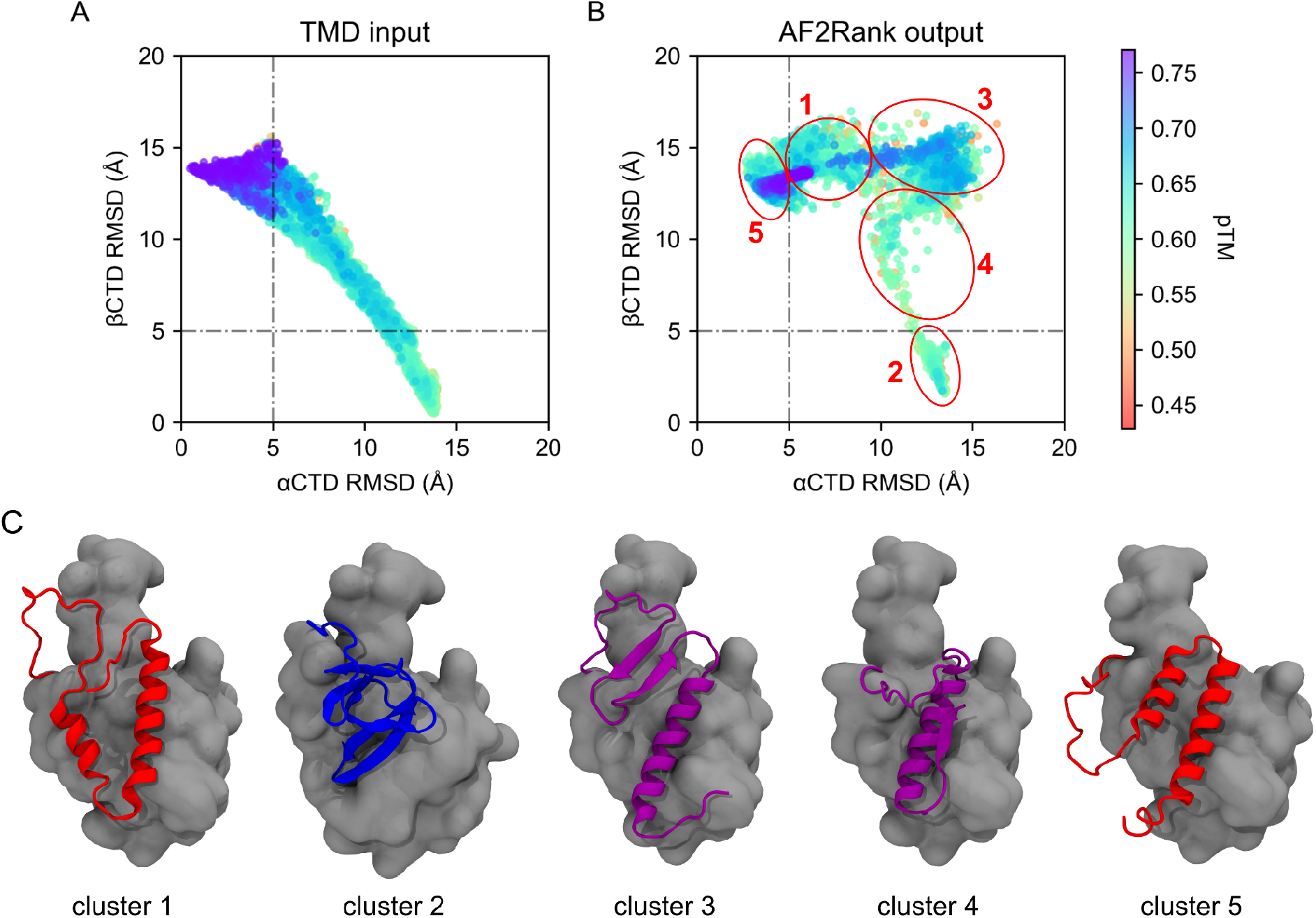
The structural diversity of RfaH is mirrored by fold-switching simulations of *E. coli* RfaH. (**A**) 2D projection of the RMSD and pTM calculated by AF2Rank for the input decoy structures obtained from nearly 200 TMD trajectories starting from either the autoinhibited or active state of RfaH. (**B**) 2D projection of the RMSD of the output structures refined by AF2Rank from the input TMD decoy structures, using the minimized structure of the autoinhibited and active states of *E. coli* RfaH. Each circle and number indicates the location around which a structural cluster was calculated. (**C**) Cartoon representation of the CTD structures for each cluster representative from the 5 relevant groups indicated in (**B**), with the NTD represented in surface representation in gray.

The AMBER-minimized autoinhibited and active states of *E. coli* RfaH were ranked favorably by AF2Rank, with pTM scores of up to 0.76 and 0.58, respectively, whereas the decoys obtained from the fold-switching simulations showed similar or lower pTM scores when compared to αRfaH or βRfaH, independent of the fold-switching TMD simulations being steered from the autoinhibited to the active state or vice versa (Figure 4A and Appendix Figure S5).

After cycling each structure once through the AF2 network, the output structures were analyzed based on their structural similarity to either the αCTD or βCTD (Figure 4B). The distribution of these structures changes dramatically after being passed through AF2, with structural clustering analysis revealing 5 distinct groups: (i) cluster 1 (2,943 structures) corresponds to autoinhibited RfaH with an upside-down αCTD; (ii) cluster 2 (1,704 structures) - to active RfaH with the correctly folded βCTD; (iii) cluster 3 (1,380 structures) - to RfaH with the previously identified mixed α/β CTD fold in which β-strands β1-β2 are forming a β-sheet, whereas helix α2 is interacting with the NTD; (iv) cluster 4 (153 structures) - RfaH with a mixed α/β CTD fold in which only helix α2 and β-strand β-5 are folded, with helix α2 interacting with the NTD; and (v) cluster 5 (1,000 structures) - autoinhibited RfaH with the correctly oriented αCTD (Figure 4C).

We note that RfaH conformations with incorrectly oriented αCTDs have been already reported by us in 19 out of 100 simulations of *E. coli* RfaH using physical- and knowledge-based potentials going from the unfolded state to the autoinhibited αRfaH state (Galaz-Davison *et al*, 2021), whereas structures of *E. coli* RfaH with a mixed α/β CTD fold have been observed in AF2 prediction of alanine scanning mutants of interdomain CTD residues, some of which were shown to be constitutively (i.e., independently of *ops*) active (Tabilo-Agurto *et al*, 2025).

By matching input TMD decoys with AF2Rank output clusters (Appendix Figure S7), it was possible to determine that the mixed α/β CTD in cluster 3 originates from input decoy structures that are dissimilar in terms of RMSD to both native CTDs and mostly obtained from TMD simulations of the fold-switch of RfaH from βCTD to αCTD, suggesting either disorder or partial β-folding. In contrast, the input decoys that do not match both native CTDs obtained from TMD simulations of the transition from αRfaH to βRfaH originate most of the structures with an upside down α-helical hairpin in cluster 1. Therefore, it can be concluded that TMD-derived structures deviating from αCTD and βCTD can be optimized by the structural module of AF2 into both CTD topologies.

When using an MSA instead, as in the case of the structure predictions of the RfaH orthologs from the different databases, a prediction of a mixed α/β CTD implies that the AF2 prediction is retrieving some signals related to the interdomain interactions with the NTD, making the distograms generated by the EvoFormer more diffuse (Appendix Figure S8), thus causing the structural module to minimize RfaH into the mixed α/β CTD conformation.

### *In vivo* transcription-translation assays of RfaH homologs matches AF2 predictions

*E. coli* RfaH controls gene expression through contacts to RNAP via its NTD and the ribosome via its βCTD. Despite sharing only 21% sequence identity, RfaH and NusG make similar contacts to both macromolecular complexes (Kang *et al*, 2018; Wang *et al*, 2020b; Molodtsov *et al*, 2024) and RfaH orthologs from diverse bacteria, including the distant RfaH homolog from *V. cholerae* (43% sequence identity to *E. coli* RfaH), act similarly to *E. coli* RfaH in a heterologous *E. coli* system *in vitro* and complement *rfaH* deletion *in vivo* (Carter *et al*, 2004).

Thus, we selected candidates with either metamorphic, mixed α/β. or monomorphic CTD from the Genomic Cluster of RfaH homologs (Appendix Table S1), which were curated using HMMs of sequence conservation and Markov clustering of genome sequences (Wang *et al*, 2020a), for functional analysis using *in vivo* reporter assays. These RfaH homologs were identified based both on their sequence conservation and on their presence in highly diverse gene neighborhood contexts (Wang *et al*, 2020a), and thus are more likely to retain canonical RfaH-like regulatory functions, as compared to those proteins identified solely by sequence conservation (e.g., from ColabFoldDB or InterPro), despite their divergent sequences.

Here, we used a well-established *in vivo* luciferase reporter assay (Appendix Table S2), in which one plasmid encodes the *Photorhabdus luminescens luxCDABE* operon positioned downstream of the *P_BAD_* promoter, while a second compatible plasmid expresses RfaH (or its variant) from an IPTG-inducible *P_trc_* promoter (Belogurov *et al*, 2010). Both plasmids are co-transformed into an *E. coli* strain lacking chromosomally encoded RfaH (DH5ɑ Δ*rfaH*). The *in vivo* luminescence activity levels are normalized by cell counts, ascertained by optical density at 600 nm (OD_600_, Appendix Figure S9).

The regions upstream of the *lux* operon can be designed to test different features of RfaH activity (Figure 5A). For example, the *lux* expression depends on RfaH when the RBS is absent, requiring ribosome recruitment mediated by the βCTD (Burmann *et al*, 2012). Expression also relies on the *ops* element when metamorphic RfaH is used because fold-switching activation of *E. coli* RfaH is triggered by interactions of RfaH with *ops*-paused RNAP (Burmann *et al*, 2012). The absence of both a functional *ops* and an RBS leads to very low levels of *lux* expression in the presence of *E. coli* RfaH, comparable to an empty vector lacking RfaH (Figure 5E and 5F), as RfaH fails to load and fold-switch, opening a path to NusG/Rho-mediated transcriptional termination (Cardinale *et al*, 2008; Lawson *et al*, 2018).

**Figure 5.**
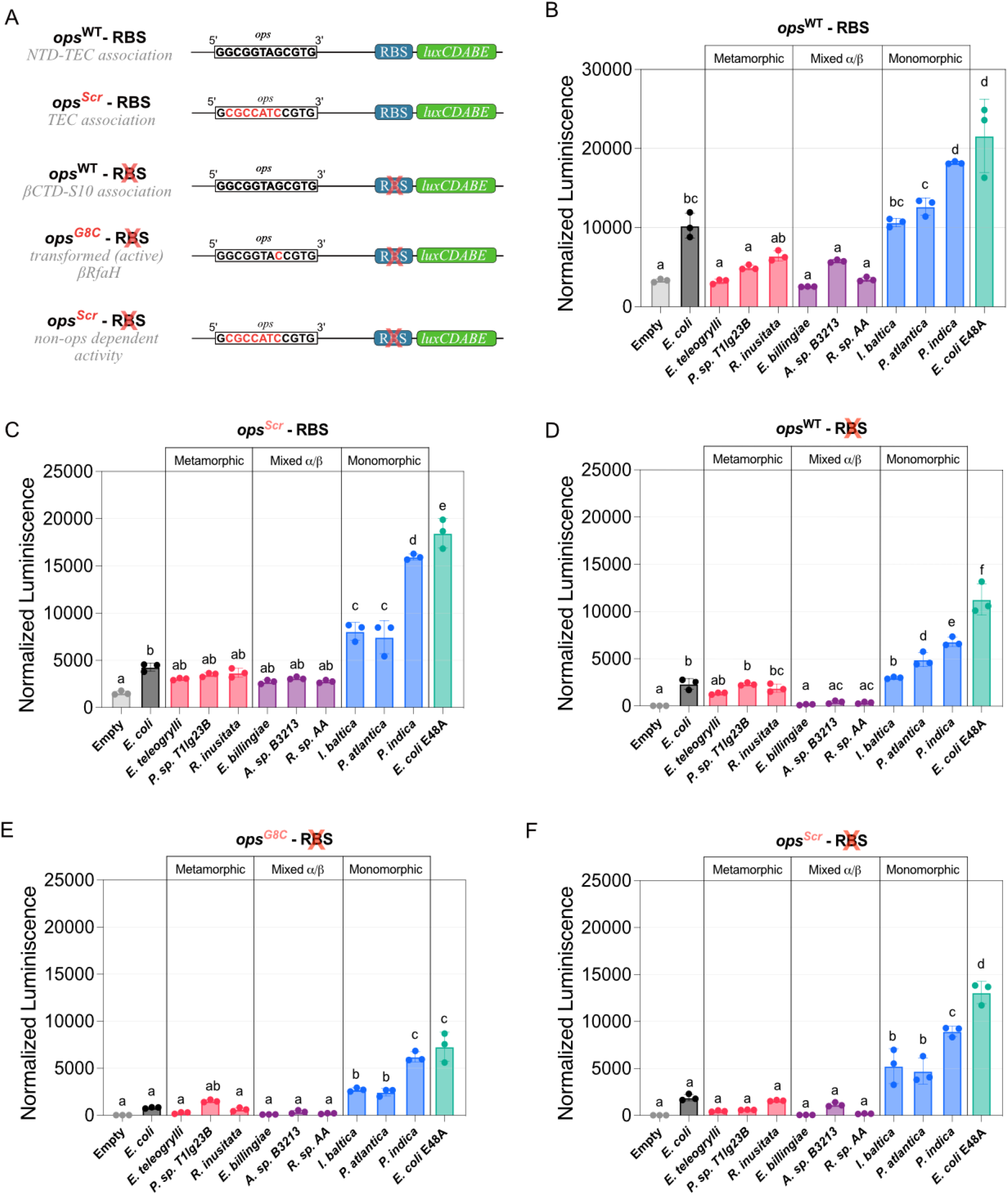
Cellular assays of RfaH homologs’ effects on gene expression. (**A**) Reporter constructs used for exploring different aspects of RfaH action on transcription-translation coupling in *E. coli* DH5α Δ*rfaH* cells. The *in vivo* activity is determined by the levels of luminescence generated by the *lux* operon in response to RfaH-regulated expression and the features of the *ops* and the presence/absence of RBS upstream of the *luxC* gene: wild-type *ops* (*ops*^WT^) and RBS (**B**), scrambled *ops* (*ops*^Scr^) and RBS (**C**), *ops*^WT^ and absent RBS (**D**), G8C mutant of *ops* (*ops*^G8C^) and absent RBS (**E**), and *ops*^Scr^ and absent RBS (**F**). Three controls per group were used in these assays: P*trc* plasmids encoding WT *E. coli* RfaH, its constitutively active E48A mutant, and an empty vector. Each bar represents the average raw luminescence level of six technical replicates of three colonies, normalized by cell density. The error bars correspond to the standard deviation of the three independent colonies (n=3). A one-way Analysis of Variance (ANOVA) was used to assess statistical significance, with all samples in an assay with a given luminescence construct compared to each other. RfaH homologs sharing the same letter are not significantly different (p > 0.05), whereas groups with different letters are significantly different (p < 0.05). The exact p-values for each plot using one-way ANOVA are provided in Appendix Tables S3-S7.

Conversely, RfaH variants with destabilized NTD-αCTD interactions, such as E48A and L142A, exhibit higher activity independent of the *ops* sequence (Burmann *et al*, 2012; Tabilo-Agurto *et al*, 2025), as demonstrated by the use of *ops* variants that induce RNAP pausing but fail to recruit RfaH, such as a scrambled *ops* sequence (*ops*^Scr^) (Belogurov *et al*, 2007) and the *ops* G8C mutant (*ops*^G8C^) (Svetlov *et al*, 2018). While autoinhibited RfaH can only be loaded at the *ops*^WT^, constitutively active variants have multiple opportunities to engage RNAP throughout transcription (Burmann *et al*, 2012; Tabilo-Agurto *et al*, 2025).

The assay results suggest that putative metamorphic orthologs exhibit *E. coli* RfaH-like activity primarily in the *ops*^WT^-no RBS context (Figure 5D). Among these, only *Erwinia billingiae* RfaH displayed intermediate activity levels between the empty vector and *E. coli* RfaH. Notably, no activity was observed with non-canonical *ops* sequences (Figures 5E and 5F), further supporting the requirement of the *ops* signal to trigger domain dissociation and the transition to the active βCTD state. The efficiency of these orthologs appears limited: under leaky, low-level expression, statistical analysis showed no significant difference compared to the control, even when both *ops*^WT^ and RBS are present (Figure 5B). However, an increase in expression upon the addition of IPTG restored their activity in the *ops*^WT^+RBS context (Appendix Figure S10), suggesting that the low activity under more stringent conditions is attributable to suboptimal recruitment or impaired fold-switching due to lower binding affinity to RNAP in the heterologous *E. coli* system.

The orthologs classified as monomorphic, and thus expected to be constitutively active, exhibited activity across all tested *ops* contexts, including in the absence of RBS (Figure 5B-F). Notably, their activity levels were comparable to the *ops*-independent *E. coli* RfaH E48A mutant, in which the key salt bridge at the NTD:CTD interface is disrupted. All monomorphic orthologs were found to lack a few key residues that make contacts to *ops* DNA in RfaH, as well as the conserved residues in loop 2 that mediate contacts with Rho in NusG (Appendix Figure S11). This mosaic architecture enables these orthologs to promote the expression of the long *lux* operon even when the RBS is absent. Our study provides the first indirect evidence of putative, constitutively active RfaH proteins, which are predicted to have the monomorphic NusG-like fold, in other bacteria.

Finally, the mixed α/β topology likely reflects fold instability, resulting in lower activity than *E. coli* RfaH in RfaH:S10-dependent translation. Indeed, this group of orthologs showed reduced activity, particularly upon removal of the RBS (Figures 5D-F), a condition where stable βCTD folding is essential for functional coupling. However, this observed lack of activity may also be attributed to expression or solubility issues, as even under conditions that induce protein expression, their activity recovery in the *ops*^WT^ with RBS architecture was less pronounced than that of their metamorphic counterparts (Appendix Figure S10).

### AF2 classification relates to phylogeny and genome location of RfaH-dependent operons

Observed *in vivo* differences in RfaH activity matched their predicted CTD folding, prompting us to explore the phylogeny of these sequences. Since AF2 uses coevolutionary information from the MSA, which is generated by pairing homologous sequences close to the input query, some evolutionary information is already incorporated into structure prediction. This is because the MSA guides the predictions via the Evoformer; therefore, alignments including different homologs would result in sampling different parts of the conformational space for the query sequence, as reported in previous works (Wayment-Steele *et al*, 2024; Lee *et al*, 2025).

As expected from the large-scale structure predictions herein and the experimental validations using a subset of RfaH orthologs, we generated a maximum likelihood phylogenetic tree using a MSA of the proteins from the Genomic Cluster using IQ-TREE 2 (Minh *et al*, 2020), which shows that similar structural predictions are shared among related members of distinct clades, particularly resulting in large clades for most of the metamorphic predictions, and smaller clades containing the mixed α/β and monomorphic members (Figure 6).

**Figure 6.**
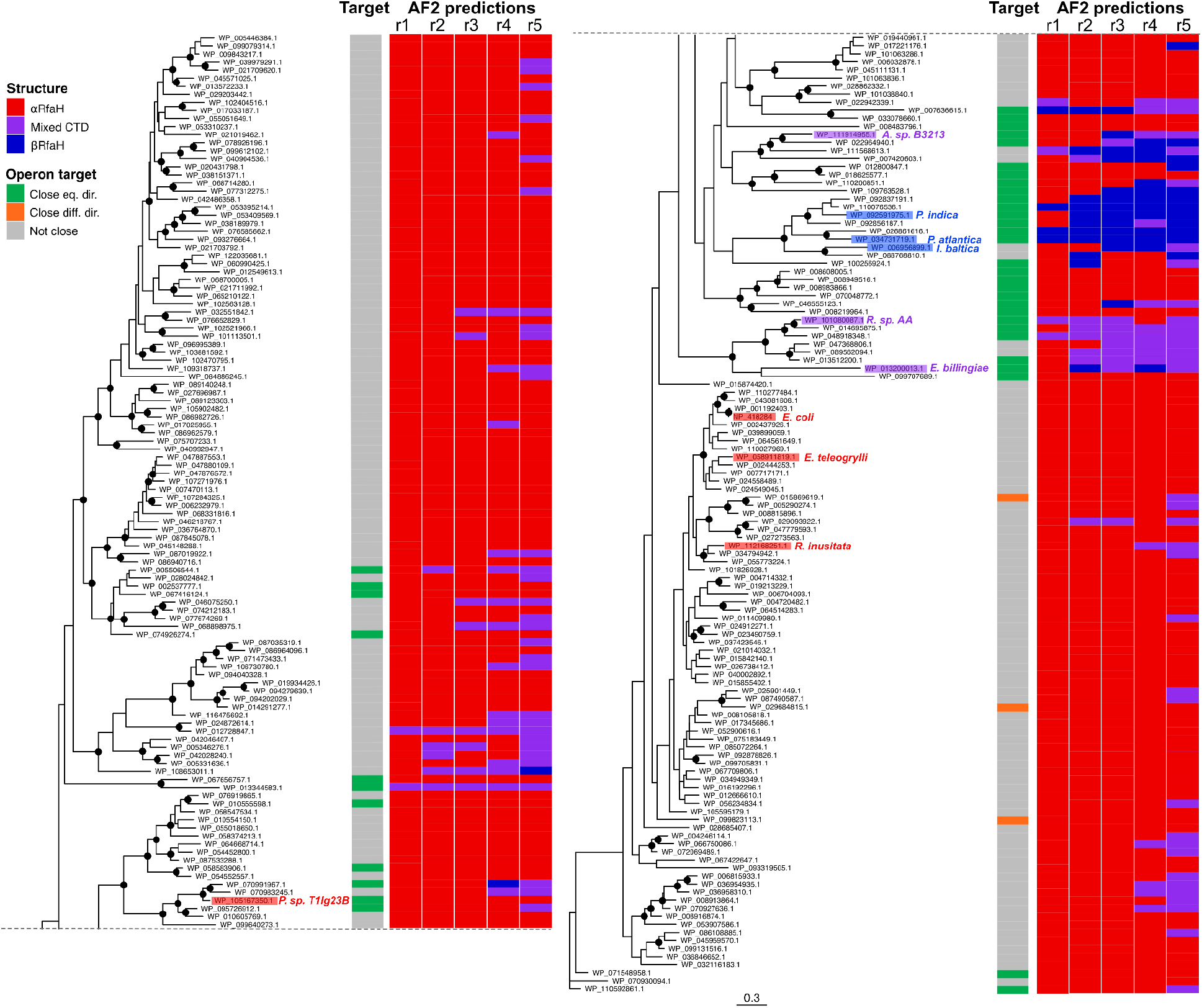
AF2 predictions are grouped by phylogeny and match long operon vicinity. RfaH-like sequences within the Genomic Cluster were used to infer a phylogenetic tree, using *E. coli* NusG as the outgroup. Nodes with Shimodaira–Hasegawa-like approximate likelihood ratio test (SH-aLRT) ≥ 80% and ultrafast bootstrap (UFBoot) ≥ 95% support are highlighted with black circles. The potential target operon of each RfaH ortholog was identified and classified as either distant (grey) or near the *rfaH* gene, and if so, whether it was in the same (green) or different (orange) direction (see Methods). The 5 models obtained for each RfaH ortholog used to construct the tree from the AF2 predictions are ranked from best (r1) to worst (r5) and color coded as metamorphic (red), mixed α/β CTD (purple) or monomorphic (blue).

The uncontrolled *ops*-independent recruitment of monomorphic RfaH could pose a challenge for the survival of the host bacterium because RfaH binds to RNAP with higher affinity than NusG (Kang *et al*, 2018) and inhibits Rho-mediated termination (Sevostyanova *et al*, 2011), which is an essential function of *E. coli* NusG (Cardinale *et al*, 2008; Lawson *et al*, 2018). A previously proposed ancestral recruitment mechanism, wherein the RfaH NTD is recruited to the transcribing RNAP while the CTD is being made by the ribosome (Belogurov *et al*, 2009), could explain how RfaH could be “selectively” targeted to avoid the lethal NusG exclusion. Since this mechanism requires that RfaH acts *in cis*, we investigated operons located in the vicinity of the *rfaH* gene within the genomes of these organisms (see *Methods*).

By tracking the relative location of *rfaH* with respect to neighbouring 30 genes and their operon organization, it can be seen that the operons surrounding genes of most metamorphic members are not adjacent to the *rfaH* gene and thus are likely not regulated by RfaH (Figure 6). This pattern is expected because *E. coli*-like metamorphic RfaH proteins have evolved to act on their targets *in trans*, by becoming activated only upon binding to their cognate (*ops*-like) target (Zuber *et al*, 2018). In that case, autoinhibition ensures that RfaH is able to control far-away operons but does not interfere with the constitutive and essential function of NusG function.

A different pattern is observed in several of the species with predicted RfaH structures with mixed α/β CTDs and in most of the monomorphic RfaH clade organisms with canonical NusG-like states, for which a long operon is located in the immediate genomic neighborhood or contains an *rfaH* gene (Appendix Figure S12). These neighbor operons in monomorphic proteins frequently encode LPS biosynthesis machinery, the known targets of RfaH (Table 1).

**Table 1.** Lipopolysaccharide-related gene in operon next to the different RfaH orthologs. A summary of genomic contexts of RfaH orthologs classified according to the AF2 predictions. The numbers indicate how many *rfaH* genes are next to an operon and whether the operon contains LPS biosynthesis genes, and the percentages next to them display the relation to the previous category, i.e, “Next to operon”/”Total” and “LPS in operon”/”Next to operon”.

| ***Category*** | **Total** | **Next to operon** | **LPS in operon** |
|---|---|---|---|
| Metamorphic | 212 | 17 (8%) | 15 (88%) |
| Mixed α/β | 12 | 8 (66%) | 1 (12%) |
| Monomorphic | 10 | 7 (70%) | 7 (100%) |

Notably, by analyzing RfaH from the Interpro sequences after matching their Genbank IDs using the UniProt ID mapping tool (UniProt Consortium, 2025), five monomorphic clades were observed in this second dendrogram (Appendix Figure S13). Among these clades, several sequences were unavailable to perform the genomic context analysis (Appendix Table S8), such as clade 1 that had no sequence available. Clade 2 was found to be composed mostly of halophilic gammaproteobacteria, containing five RfaH sequences, of which three were close to a long operon, yet none of those contained LPS-related sequences. Clade 3 contained the RfaH from *P. indica*, already analyzed above, while clade 4 contained eight Pseudomonadaceae bacteria, three of which were close to a target operon, only one containing LPS-related genes.

Finally, clade 5 contained primarily lithotrophic gammaproteobacteria, with two out of the six found displaying RfaH close to target operon, none of which contained LPS-related genes. Overall, these results show that monomorphic RfaHs are generally closer to target operons than metamorphic RfaHs. This is consistent with a *cis* regulatory mechanism, in which these monomorphic RfaHs that bind and regulate the RNAP function in an *ops*-independent manner, control the target operon transcription by being expressed right next to them. If these monomorphic proteins are either rapidly degraded or aggregate due to their hydrophobicity, this *cis* mechanism could explain how these proteins are able to regulate RNAP without interfering with the essential function of NusG in their host organism.

### Experimental validation of a predicted monomorphic RfaH ortholog by circular dichroism

To experimentally test the AF2 predictions of a monomorphic RfaH ortholog, we were able to purify (Appendix Figure S14) and collect circular dichroism (CD) spectra of the putative monomorphic *P. atlantica* RfaH, *E. coli* RfaH, and *E. coli* NusG as a control. As expected, the CD spectrum of *P. atlantica* and *E. coli* RfaH differed, with the former lying above those of NusG and *E. coli* RfaH. While all three spectra display the characteristic minimum at ∼208 nm, both *P. atlantica* RfaH and NusG lack the second minimum at ∼220 nm that is characteristic of predominantly α-helical proteins (Figure 7A). Notably, *P. atlantica* RfaH exhibited the lowest estimated α-helical content (13%) as compared to NusG (14%) and *E. coli* RfaH (17%), and the highest β-strand content (27%) compared to NusG (25%) and *E. coli* RfaH (21%) (Figure 7B), largely consistent with their solved and predicted structures (Appendix Figure S15). Together, these spectral features and the CD-derived secondary structure content further support the classification of *P. atlantica* RfaH as a monomorphic ortholog.

**Figure 7.**
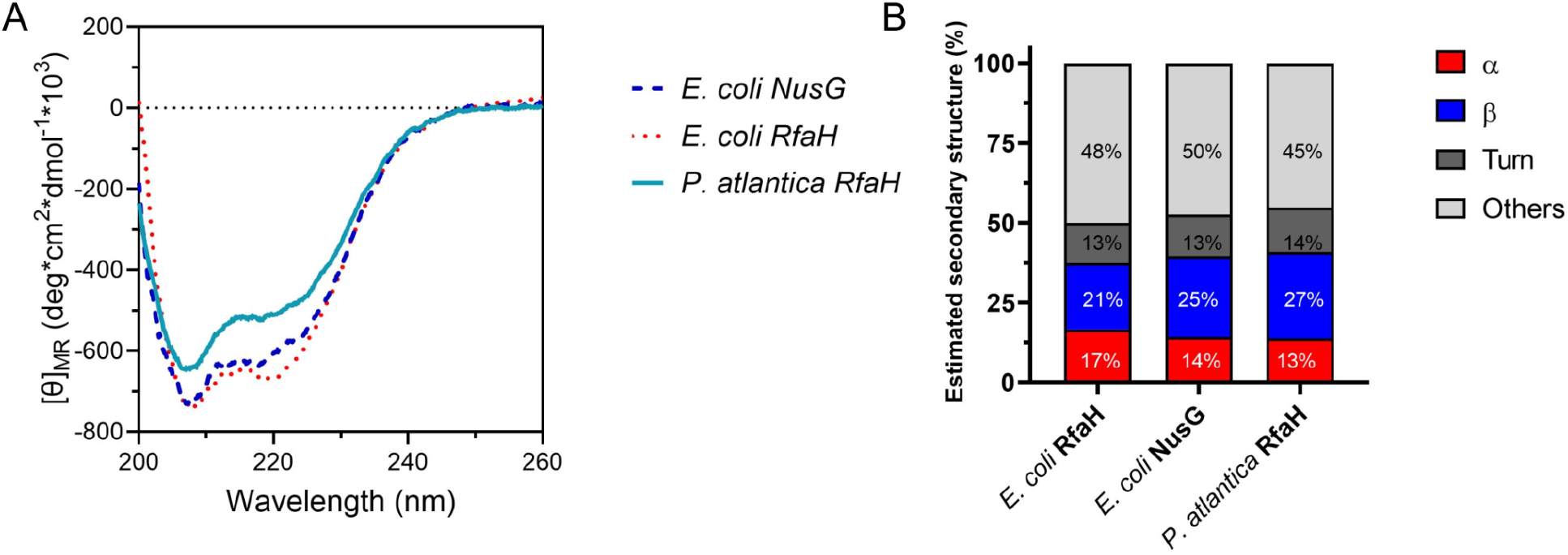
Experimental validation of a monomorphic RfaH ortholog. (A) Far-UV CD spectra, expressed as mean residue molar ellipticity, of *P. atlantica* RfaH, *E. coli* RfaH, and *E. coli* NusG (control). (B) Helix (regular helix + distorted helix) and strand (parallel + antiparallel) content estimated from the CD spectra in (A) using BeStSel (Micsonai *et al*, 2025).

## DISCUSSION

Metamorphic proteins present a challenge for DL structure prediction algorithms as the predominant fold is commonly observed. AF2 in its default implementation is not exempt from this issue (Chakravarty & Porter, 2022), prompting the use of different approaches to discover alternate structures, such as sequence-based (Wayment-Steele *et al*, 2024) and energy-based MSA clustering (Guan *et al*, 2024) or random sampling approaches (Lee *et al*, 2025). An alternative approach, as utilized herein, is to cluster the sequences from a given protein family, predict their three-dimensional structures, and analyze their structural differences. If the MSA is representative of the orthologs in the family, so that each clade member can be predicted accurately, then the structural differences therein are likely arising from changes in sequence conservation and coevolution.

In this work, we used RfaH, a metamorphic ubiquitous transcription factor for which a wealth of functional (Belogurov *et al*, 2010; Burmann *et al*, 2012; Shi *et al*, 2017; Tabilo-Agurto *et al*, 2025), structural (Burmann *et al*, 2012; Zuber *et al*, 2018; Kang *et al*, 2018; Zuber *et al*, 2019, 2024), and *in silico* modeling (Wayment-Steele *et al*, 2024; Tabilo-Agurto *et al*, 2025; Lee *et al*, 2025) and simulation data (Ramírez-Sarmiento *et al*, 2015; Gc *et al*, 2015; Seifi *et al*, 2021; Galaz-Davison *et al*, 2021; Appadurai *et al*, 2021; Zuber *et al*, 2024; González-Higueras *et al*, 2024) are available to evaluate the potential of this approach. Our present results argue that the prediction of monomorphic and mixed secondary structures of fold-switching CTD in the RfaH family does not necessarily represent failures in AF2 predictions. Instead, we posit that these “unexpected” folds correspond to RfaH orthologs that lack a fold-switching mechanism of fully-evolved RfaH and assume the constitutive active state. Our *in vivo* activity assays, phylogenetic analysis, and examination of genomic neighborhoods support the existence of previously hypothesized monomorphic RfaH orthologs which could be recruited to their targets in *cis*. While limited solubility precluded analysis of other putative monomorphic RfaH orthologs, our CD data show that at least one such candidate, *P. atlantica* RfaH, has an apparent secondary structure content that is more similar to NusG than *E. coli* RfaH.

To the best of our knowledge, this is the first report to describe putative monomorphic RfaH orthologs with distinct functional patterns, diverging from the known metamorphic RfaH. Based on our findings, we propose a new evolutionary branch within the RfaH family, in which some orthologs have bypassed the metamorphic requirement to achieve constitutive activity, likely as an adaptation to specific genomic and regulatory contexts (Figure 8).

**Figure 8.**
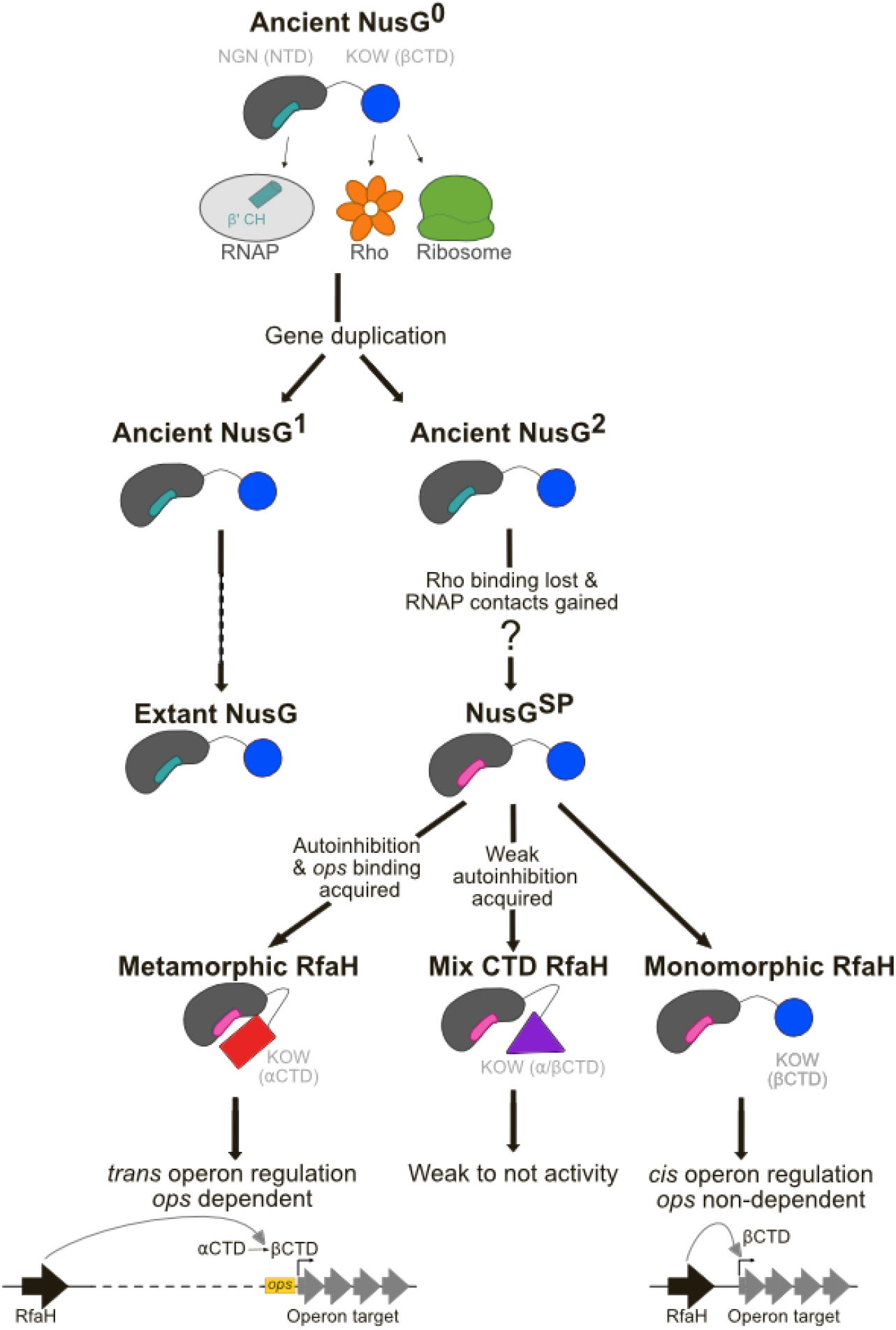
Step-wise structural and functional path from ancient to extant RfaH orthologs.

Our findings are in line with the previous hypothesis regarding the emergence of RfaH within the universally conserved family of NusG transcription factors (Belogurov *et al*, 2009). Under that model, a gene duplication event produced an intermediate variant (NusG^2^ in Figure 8) that lost its Rho-binding capability and acquired a deletion in the NTD that reduced the protein’s overall size and remodeled its hydrophobic profile. Crucially, this intermediate retained an exposed, hydrophobic RNAP-binding region, a feature shared by monomorphic RfaH and the ancestor of all RfaH orthologs (NusG^SP^ in Figure 8). This increased hydrophobicity would have reduced solubility, restricting its regulatory activity to the site of synthesis, i.e. in *cis*. Indeed, when structural alignment is used to identify conserved NTD residues that bind to RNAP, orthologs contain more than 70% hydrophobic residues (Appendix Figure S11) at those positions. This percentage is much closer to that of RfaH (80%) than NusG (57.14%). The protein only regains solubility and the ability to operate *in trans* when its CTD refolds into a helical conformation, burying those very residues. Ultimately, our results strengthen this evolutionary model by demonstrating that several extant RfaH orthologs appear to resemble this insoluble, *cis*-acting ancestral state.

Previous reports suggested that RfaH family sequences with non-metamorphic features were likely from its non-switching paralog NusG (Galaz-Davison *et al*, 2022; Porter *et al*, 2022). RfaH and NusG possess a mixture of parallel (transcription processivity and coupling to translation) and orthogonal (Rho-dependent transcription termination) functions (Wang & Artsimovitch, 2020), and sequence alone is insufficient to discern the homology relationship within this family. Here, we focused on RfaH homologs that were clustered using HMMs alongside *E. coli* RfaH and expected to be metamorphic (Belogurov *et al*, 2009; Wang *et al*, 2020a). However, our results suggest that some members of this group are monomorphic, and their genomic context is different from those of both the canonical metamorphic RfaH and the monomorphic NusG. Furthermore, when extending our genomic context analysis to five monomorphic RfaH clades from the Interpro dataset, we observed roughly a third to half of the sequences had RfaH next or inside a long operon, with one of those containing one gene related to LPS synthesis. We also find that, like RfaH, its monomorphic homologs activate expression of a horizontally-acquired *P. luminescens lux* operon. This activity contrasts with the canonical function of NusG, which aids Rho in termination transcription of foreign DNA (Lawson *et al*, 2018). Thus, the putative monomorphic RfaH homologs act as RfaH and not as NusG.

RfaH promotes gene expression by counteracting Rho/NusG silencing through many mechanisms, all of which require RfaH activation through the *ops*-mediated fold switch (Wang & Artsimovitch, 2020). The requirement for *ops* can be bypassed by weakening the NTD-CTD interactions to stabilize the active (monomorphic) state, as observed with the constitutively active (*ops*-independent) E48A RfaH variant (Burmann *et al*, 2012). We show that the putative monomorphic RfaH homologs activate *lux* expression similarly to E48A (Figure 5) and are encoded near their putative target operons (Figure 6), supporting the idea that these proteins are not autoinhibited and achieve selective recruitment *in cis*.

It is worth noting that, in the absence of RBS (Figure 5C-F), the putative monomorphic RfaH orthologs have similar or lower *in vivo* activity than the *E. coli* RfaH E48A mutant; a similar mutant (E48S) exhibits a 1:1 equilibrium between the autoinhibited and active states (Burmann *et al*, 2012). This observation can be partly explained by two factors. First, sequence divergence and expression levels may limit functional compatibility with the host machinery. Highly divergent *V. cholerae* RfaH ortholog, which shares only 43% sequence identity with *E. coli* RfaH but is predicted to fold into the autoinhibited state (Artsimovitch & Ramírez-Sarmiento, 2022), maintains both *ops*-dependent recruitment and hemolysin secretion in the Δ*rfaH E. coli* strain, yet exhibits transcription elongation defects *in vitro*, requiring a 5-fold higher concentration than *E. coli* RfaH to match increased elongation rates of *E. coli* RNAP (Carter *et al*, 2004). Low *in vivo* protein titers or structural mismatches between the monomorphic orthologs and *E. coli* RNAP may prevent higher luciferase expression relative to the E48A mutant. This limitation is supported by the fact that IPTG-induced overexpression rescues activity when an RBS is present (Figure 5B). Second, these proteins may be predominantly folded in the active state while still transiently populating the autoinhibited state. Furthermore, the three monomorphic proteins were overexpressed and purified using a His-tag, yet only one of them displayed sufficient solubility to perform circular dichroism assays, revealing it to contain similar helical and extended secondary structure content to NusG, and different to that of *E. coli* RfaH. This evidence supports the notion of extant monomorphic RfaH orthologs that are mostly, if not completely, populating the active state of RfaH (Porter *et al*, 2022).

Another predicted state with mixed α/β CTD structure was observed as part of the AF2 predictions of RfaH homologs. Our analysis of the transition between αRfaH and βRfaH for *E. coli* RfaH through TMD simulations, which were then input to AF2Rank, revealed the presence of five well-defined structural clusters, two of which correspond to its autoinhibited and active states, and three clusters representing alternative conformations: (i) RfaH with an upside-down αCTD, RfaH with only helix α2 and β-strand β5 folded and, importantly, the mixed α/β state observed in 9% of the predicted structures of RfaH homologs. Before passing through AF2Rank, the structures exploring the transition between native states display a high RMSD to both experimental structures, likely exploring high energy parts of the RfaH free energy landscape, as shown previously (Galaz-Davison *et al*, 2021).

Coincidentally, after passing through the AF2 architecture, the RfaH structures with mixed α/β CTD remain at the high energy part of the landscape and display lower pTM and lower mean pLDDT than proteins with αRfaH and βRfaH folds. Although these confidence metrics are not directly related with experimental stability, together with low *in vivo* activity of the mixed α/β CTD variants without RBS (Figure 5), they are consistent with an unstable fold. From an evolutionary standpoint, the instability caused by the presence of an exposed NTD can be interpreted as a signal of structural ambiguity in protein sequences relating to emerging folds and fold-switching (Tian & Best, 2020), suggesting that mixed α/β orthologs are still undergoing structural and functional diversification towards *E. coli*-like, fully evolved RfaH. Interestingly, a recent nuclear magnetic resonance spectroscopy study of the *E. coli* RfaH CTD, aimed to uncover transient states potentially en route of the αCTD interconversion (Cai *et al*, 2025), described an intermediate state (populated in ∼23% of the captured ensembles) in which a β-hairpin formed by β-strands β1-β2 lies on top of a transient α-helix α5 that corresponds to helix α2 in our article. This finding is fully consistent with the mixed α/β CTD structures found both in our TMD simulations and our AF2 predictions of divergent RfaH orthologs, highlighting the structural feasibility of this emerging fold, and its likelihood as a result of evolutionary transitions towards autoinhibited RfaH.

Phylogenetic analysis reveals that AF2 predictions with similar fold propensities are clustered in well-defined clades. Although AF2 and phylogenetic inference are separate experiments, a correlation between them is expected for structural predictions that are based on evolutionary information. This correlation implies that the sequence variability was sufficient to detect the mixed α/β and monomorphic RfaH homologs in the AF2 protein structure predictions, which have different genomic contexts, *in vivo* activity profiles and estimated secondary structure content, supporting the idea that their structural diversity is associated with distinct functional outcomes.

## METHODS

### Large-scale protein structure prediction

Amino acid sequences for *E. coli* RfaH homologs were retrieved from three databases: (a) ColabFoldDB (Mirdita *et al*, 2022) (468 sequences), accessed on January 16 2026 via MMseqs2 (Steinegger & Söding, 2017) using the sequence of *E. coli* RfaH (UniProt accession ID P0AFW0) as input for ColabFold v1.5.5 (Mirdita *et al*, 2022) and filtered based on a minimum sequence coverage of 95%, a bit-score of 140 and a maximum e-value of 1e-30; (b) InterPro (Blum *et al*, 2025) (family entry IPR010215, 3,058 sequences), accessed on January 3 2024; and (c) a previously reported list of 416 RfaH sequences (Cluster 1) curated by HMMs with known genomic context that were classified by Markov clustering and include *E. coli* RfaH (Genomic Cluster, hereafter) (Wang *et al*, 2020a). In the case of the ColabFoldDB sequences, these were filtered to eliminate duplicate sequences between this database and InterPro using CD-HIT v4.8.1 (Fu *et al*, 2012), and *E. coli* RfaH (UniProt accession ID P0AFW0) was manually added.

These 3,942 RfaH-like sequences were used as input for DL-guided protein structure prediction using a local version of ColabFold v1.5.5 (Mirdita *et al*, 2022) (LocalColabFold, available at https://github.com/YoshitakaMo/localcolabfold) using default parameters: 1 seed, 5 models, alphafold2_ptm model for monomers, 3 recycles, no templates, MSAs generated using MMseqs2 (Steinegger & Söding, 2017). The resulting 19,710 structures were analyzed using TM-align v20220412 (Zhang & Skolnick, 2005) to calculate the TM-score (Zhang & Skolnick, 2004), a metric to assess the similarity between two protein structures ranging from 0 (no match) to 1 (exact match), and the root mean square deviation (RMSD) in Å for the full-length protein and their CTDs, using the best ranked predicted structure of *E. coli* αRfaH and its αCTD and the experimental structure of the βCTD of *E. coli* (PDB ID 2LCL) as references. The per-residue secondary structure of the CTD of the predicted structures was analyzed using STRIDE v1.6.4 (Frishman & Argos, 1995). ColabFold was also used to obtain the distograms (set at a 1 nm cutoff) of *E. coli* RfaH in the active and autoinhibited states, obtained using 0 and 3 recycles, respectively, as well as the distograms of the 9 selected RfaH orthologs from the Genomic Cluster for experimental characterization after 3 recycles.

Frequency histograms of the percentage of α-helices (classified as ‘H’ in STRIDE) and β-strand (classified as ‘E’ in STRIDE) for all predicted structures from all databases were first used to determine potential cutoffs to distinguish between metamorphic αRfaH, monomorphic βRfaH and RfaH with mixed α/β CTD (Appendix Figure S1). Based on this analysis and the histograms in the plots of α-helical and β-strand secondary structure, metamorphic αRfaH were classified based on having a percentage of α-helices > 32.5% and < 2.5% of β-strands, monomorphic βRfaH on having a percentage of β-strands > 30.0% and < 2.5% of α-helices, and RfaH with mixed secondary structure in their CTD on having a percentage of α-helices and β-strands > 2.5%. Other secondary structure ranges were classified as ‘none’. To determine that these values were adequate for our analysis, we performed a sensitivity test (Appendix Figure S16) by changing the thresholds by ±5% over the data for all structures predicted from all databases, showing that the predictions of RfaH orthologs with monomorphic CTD and mixed secondary structure in their CTD is robust, and only metamorphic RfaH orthologs were reduced with an increase in uncategorized structures (Appendix Figure S16). The data was further manually curated based on visualization of the predicted structures, which are presented throughout the manuscript through images generated using VMD v1.9.3 (Humphrey *et al*, 1996).

Additionally, salt-bridge distances were calculated between residues equivalent to the E48-R138 pair in *E. coli* RfaH for all predicted structures by determining these residues from each protein based on a MSA generated for each database, including *E. coli* RfaH, using MAFFT v7.526 (Katoh & Standley, 2013) in auto mode, and a custom python script to get the coordinates of the Cβ atoms of the sidechain of these residues (Cα in the case of glycine) and measure their distance. Lastly, sequence logos for the residues in these positions of the alignment with respect to *E. coli* RfaH were calculated from the MAFFT-generated MSAs for each database using Logomaker v0.8 (Tareen & Kinney, 2020).

### TMD simulations

The full-length structures of the autoinhibited and active states of *E. coli* RfaH were obtained from a previous work (González-Higueras *et al*, 2024). Briefly, αRfaH was generated using the experimentally solved structures of its autoinhibited state (PDB 5OND, chain B, NTD; PDB 2OUG, chain B, αCTD), whereas for βRfaH, the αCTD was replaced by the experimentally solved structure of βCTD (PDB 2LCL). Missing linker residues between the NTD and CTD were added using MODELLER v9.23 (Webb & Sali, 2016). Both models were energy minimized after being solvated in a cubic box of 1.0 nm of padding with TIP3P water molecules and neutralized with counterions using GROMACS v4.5.3 (Pronk *et al*, 2013) and the Amber ff99SB-ILDN force field (Lindorff-Larsen *et al*, 2010) using the steepest descent method.

The TMD simulations presented herein were performed similarly to a previous work from our group (Ramírez-Sarmiento *et al*, 2015), in which the collective variable that steers RfaH *E. coli* from one native state into another is defined by applying an external potential to minimize the root mean square deviation (RMSD) of the protein coordinates relative to the target final structure during the simulation trajectory, according to equation 1:

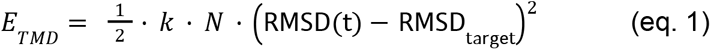

Where *k* is the force constant, *N* is the number of atoms used as a collective variable during the TMD simulation, RMSD_target_ is the target RMSD and RMSD(t) is the calculated frame at each timestep during the simulation.

The main differences with our previous work are: (i) the use of the RMSD for the backbone atoms of residues 110-162 in full-length *E. coli* RfaH as collective variable (totaling 252 atoms); and (ii) the use of an implicit solvent model, which reduces the computational cost and, consequently, allows to perform multiple TMD simulations using different external potential force constants to steer *E. coli* RfaH towards either the autoinhibited or the active state.

Specifically, we utilized tleap from AmberTools v22.3 (Case *et al*, 2023) to parameterize our simulations systems for *E. coli* αRfaH and βRfaH using the ff19SB force field (Tian *et al*, 2020) and the GB-Neck2 implicit solvent model with mbondi3 intrinsic radii to accurately represent the solvation environment (Nguyen *et al*, 2013) to perform TMD simulations using AMBER 22 (Case *et al*, 2025). First, we minimized each system for 2,500 steps using a steepest descent gradient, followed by 47,500 steps of conjugate gradient, before performing 300 short (100 ps) TMD simulations from the autoinhibited to the active state and vice versa (600 TMD simulations total) at a temperature of 298 K controlled by a Langevin thermostat and the SHAKE algorithm to constrain hydrogen-atom bond lengths to allow simulations with a time step of 2 fs, using randomly selected force constants ranging from 1.0×10^−4^ to 3×10^−2^ kcal/(mol·Å^2^) applied over the backbone atoms of residues 110-162, while aligning the protein during the simulation based on the backbone atoms of the NTD, with a target RMSD of 0 Å to the final reference structure. Frames were collected every 2 ps, thus generating 50 frames per trajectory for analysis.

To select a representative number of TMD trajectories that explored all possible configurations across the fold-switch from αRfaH to βRfaH and vice versa, a universal Principal Component Analysis (PCA) map was constructed using the 10 trajectories from each set of TMD simulations in each direction of the RfaH fold-switch that reached the minimum RMSD using cpptraj v6.4.4 from AmberTools v22.3 (Case *et al*, 2023). With this data, a coordinate covariance matrix was calculated using the Cα atoms of the fold-switching CTD (residues 110–162), while the NTD was used as a rigid-body reference for alignment to remove translational and rotational motions. The covariance matrix was diagonalized to obtain the eigenvectors, upon which all 600 trajectories were projected onto the first two principal components (PC1 and PC2) of this universal PCA map (Appendix Figure S6), allowing for a direct structural comparison of the hysteresis between the different directions of the TMD simulations.

Lastly, to isolate the most representative and structurally diverse transition pathways, *k*-means clustering was performed on the projected PCA data. For each of the 300 TMD simulation datasets, the trajectory frames were clustered into *k* = 100 distinct groups based on their Euclidean distance in the PC1/PC2 subspace. The centroid of each cluster was identified, and the specific trajectory containing the centroid frame was selected as a representative pathway. This process yielded a curated dataset of 100 TMD trajectories for the simulations from αRfaH to βRfaH (3,750 decoy structures) and 99 TMD trajectories for the simulations from βRfaH to αRfaH (3,700 decoy structures), ensuring broad sampling of the fold-switching transitions in each direction while removing redundant paths.

### Decoy analysis using AF2Rank

The original Jupyter notebook of AF2Rank for Google Colab (Roney & Ovchinnikov, 2022), available on GitHub (https://github.com/jproney/AF2Rank), was modified to analyze all 7,450 decoy structures generated by TMD using both the AMBER-minimized αRfaH and βRfaH states of *E. coli* RfaH as native states. AF2Rank was run using default parameters (1 recycle, alphafold_ptm model and 1 model per input decoy) as well as masking the sequence and sidechains of the structures, in order to prevent AF2 from using its knowledge of evolutionary couplings or specific amino acid sequences to weigh in when evaluating the physical plausibility of the decoy structures and generating the output structures after passing through the EvoFormer and Structure Module.

After obtaining the output structures from AF2Rank, they were subjected to *k*-means clustering based on the Cα RMSD of their CTDs (residues 110-162) against the CTDs of the AMBER-minimized αRfaH and βRfaH states of *E. coli* RfaH. Structures were grouped in five clusters (*k* = 5) and the medoid structure from each cluster was selected as representative structure for visualization using VMD v1.9.3 (Humphrey *et al*, 1996).

### *In vivo* luminescence assay

DH5ɑ Δ*rfaH* (IA149) was co-transformed with two plasmids. One chloramphenicol-resistant plasmid encoded one of the 9 selected RfaH orthologs from the Genomic Cluster (Appendix Table S1) with at least 2/5 AF2 models predicted in the metamorphic αRfaH state (GenBank accession ID WP_058911819.1, *Erwinia teleogrylli*; WP_105167350.1, *Pseudoalteromonas sp.* T1lg23B; WP_112168251.1, *Rahnella inusitata*), the monomorphic βRfaH state (WP_006956899.1, *Idiomarina baltica*; WP_034731719.1, *Pseudidiomarina atlantica*; WP_092591975.1, *Pseudidiomarina indica*) or with a mixed α/β CTD (WP_013200013.1, *Erwinia billingiae*; WP_111914955.1, *Aliidiomarina sp.* B3213; WP_101080081.1, *Rahnella sp.* AA), *E. coli* RfaH or constitutively active *E. coli* RfaH E48A downstream of a *P_trc_* promoter that can be used to induce protein overexpression by addition of isopropyl-β-D-thiogalactopyranoside (IPTG). The second ampicillin-resistant plasmid encoded the *luxCDABE* operon downstream of several genetic constructs of *ops* (*ops*^WT^, *ops*^G8C^, *ops*^Scr^) with and without an RBS site under the control of a *P_BAD_* promoter (Appendix Table S2).

Single colonies co-transformed with both plasmids were grown overnight (ON) at 37°C in liquid LB medium (US Biological, USA) supplemented with antibiotics (30 μg/mL chloramphenicol [US Biological, USA] and 100 μg/mL carbenicillin [Applied Biological Materials, Canada]]). The following day, saturated cultures were diluted to an initial optical density at 600 nm (OD_600_) of 0.05 in 3 mL of fresh LB medium supplemented with 0.1% glucose (Sigma-Aldrich, USA) and the same concentration of antibiotics. Cells were grown until reaching a similar late exponential phase OD_600_ ≈ 0.7-0.8 (Appendix Figure S9). For IPTG-induced conditions (Appendix Figure S10), cultures were grown to an OD_600_ of 0.2–0.3 and then induced with 0.2 mM IPTG (US Biological, USA) for 1 h. The final OD_600_ was measured using a UV/Vis spectrophotometer (Jasco V-730, Jasco Inc. Japan), and luminescence was recorded in black 96-well plates using a multi-mode plate reader (Synergy HTX, BioTek Instruments, USA). Three biological replicates were performed per RfaH protein per condition, as well as for a negative control with a chloramphenicol-resistant plasmid not encoding any RfaH protein (empty).

### Phylogenetic tree inference

The sequences from the genomic cluster were subjected to clustering using CD-HIT v4.8.1 (Fu et al, 2012) at a sequence identity cutoff of 85%. An MSA of the clustered sequences, in which *E. coli* NusG was included, was generated with MUSCLE (Edgar, 2004), available at the EMBL-EBI Job Dispatcher web server (Madeira *et al*, 2024). After several iterations of manual curation, the final alignment, comprising 231 sequences, was used to infer the phylogenetic tree by maximum likelihood using IQ-TREE 2 (Minh *et al*, 2020) with the built-in automatic best-fit evolutionary model selection, which in this case was determined to correspond to LG (Le & Gascuel, 2008) with a proportion of invariant sites and gamma-distributed rate heterogeneity across sites (LG+I+G4). Maximum-likelihood tree inference was performed with an extended search of 2,000 trees, and branch support was assessed using 2,000 replicates of the Shimodaira–Hasegawa-like approximate likelihood ratio test (SH-aLRT) and 2,000 ultrafast bootstrap (UFBoot) replicates. Support values from both methods were mapped onto the resulting maximum-likelihood phylogeny. *E. coli* NusG was used as the outgroup to root the tree.

### Genomic context analysis

RfaH homologs from the Genomic Cluster were submitted to TREND (Gumerov & Zhulin, 2020) (https://trend.evobionet.com/) to extract neighboring genes (up to 15 genes on each side). The resulting datasets were downloaded and analyzed. Gene neighbors were sorted by genomic coordinates, and we applied the following criteria to determine whether an RfaH homolog is located near its potential target operon:

Step 1 – Identification of long operons. Since RfaH is required for transcription of long operons, a long operon was first defined based on the following conditions: (i) all constituent genes have the same transcriptional direction; (ii) the intergenic distance between adjacent genes is ≤200 bp; and (iii) the total length of the operon exceeds 5,000 bp.

Step 2 – Assessment of distance between RfaH homolog and operon. If an operon was identified near an RfaH homolog, we calculated the genomic distance to determine whether they are in close proximity. Three scenarios were considered: (i) (RfaH homolog and the operon has same “+” direction) or (RfaH homolog has “-” and the operon has “+” direction). To be close to the operon, absolute value of ((the end of RfaH homolog) - (start of first gene of operon)) < 1000 bp; (ii) (RfaH homolog and operon has same “-” direction) or (RfaH homolog has “+” and operon has “-” direction). Note that in the case of an operon showing “-“ orientation, the last gene of the operon is nearest to the promoter because the genes are ordered according to coordinates. The absolute value of ((the start of RfaH homolog) - (end of last gene of operon)) < 1000 bp. (iii) If the RfaH homolog is located inside the operon, it is automatically considered adjacent to it.

### Overexpression and solubility test

Plasmids encoding RfaH orthologs classified as βCTD were transformed in *E. coli* BL21(DE3) grown from an overnight LB culture and then grown in flasks with 50 mL of TB medium supplemented with kanamycin (50 μg/mL; VWR, USA) at 37°C until reaching an optical density at 600 nm (OD_600_) between 0.6 and 0.8. Protein overexpression was induced for 2 h at 37°C and overnight at 20°C with 250 rpm agitation after the addition of 0.2 mM IPTG (US Biological, USA). Cells were harvested by centrifugation (4,200 g, 10 min, 4°C) and stored at −20°C. The next day, cells were resuspended in buffer LysT (50 mM Tris-HCl pH 6.9, 1 mM β-mercaptoethanol, 500 mM NaCl, 0.1 mM PMSF) and lysed by sonication on ice and then centrifuged (12,500 g, 30 min, 4°C). The solubility was ascertained by running a SDS-PAGE gel of the supernatant and pellet sample, as well as a control without IPTG induction. In addition, the pellet was treated with LysT supplemented with 1% CHAPS (Biotium Inc, USA). *P. atlantica* RfaH encoded in the plasmid pDP26 was the only putative monomorphic RfaH exhibiting soluble expression in these conditions (Appendix Figure S14).

### Protein purification

Plasmids encoding *E. coli* RfaH (EcoRfaH) and *P. atlantica* RfaH (pDP26) were transformed into *E. coli* BL21 (DE3). Isolated colonies were grown ON in 5 mL LB culture supplemented with kanamycin (50 μg/ml). The next day, the oversaturated culture was grown in a flask containing 500 ml of TB medium supplemented with kanamycin (50 μg/ml) at 37 °C until an OD_600_ between 0.6 and 0.8 was reached. Protein overexpression was induced ON at 20 °C with 0.2 mM IPTG and agitation at 250 rpm. Cells were harvested by centrifugation (4,500 g, 10 min, 4°C) and stored at −20°C.

Cells were resuspended in buffer LysT 6.9 (50 mM Tris-HCl pH 6.9, 1 mM β-mercaptoethanol, 500 mM NaCl, 0.1 mM PMSF) supplemented with 1% CHAPS and lysed by sonication on a QSonica Q125 sonicator (10 s ON, 20 s OFF for a total of 15 min at 40% power) on ice and then centrifuged (12,500 g, 30 min, 4°C). The supernatant was loaded onto a pre-equilibrated 1mL HisTrap HP column (Cytiva, USA) with buffer LysT supplemented with 1% CHAPS. The column was washed twice with: (1) 20 mL buffer LysT 6.9 supplemented with 1% CHAPS supplemented with 20 mM imidazole (Sigma-Aldrich, USA), and (2) 10 mL buffer LysT 6.9 supplemented with 1% CHAPS supplemented with 50 mM imidazole. Elution was performed under the same buffer conditions, but with LysT buffer supplemented with 300 mM imidazole.

The fraction recovered was diluted 3 times with buffer 6.9 A (50 mM Tris-HCl pH 6.9, 1 mM β-mercaptoethanol, 0.1 mM EDTA), and load to a pre-equilibrated 1 mL HiTrap Heparin HP column (Cytiva, USA) with 10% buffer 6.9 B (50 mM Tris-HCl pH 6.9, 1.5 M NaCl, 1 mM β-mercaptoethanol, 0.1 mM EDTA). Fractions were eluted with a linear gradient of buffer 6.9 A supplemented with 100% 6.9 B. Fractions containing the desired purified fraction, ascertained by the gel analysis, were collected, concentrated and stored at −80°C with 40% glycerol.

The plasmid encoding *E. coli* NusG (pI244) was transformed into *E. coli* BL21(DE3). Isolated colonies were grown ON in 5 mL LB culture supplemented with kanamycin (50 μg/ml). The next day, the oversaturated culture was grown in a flask containing 500 ml of LB medium supplemented with kanamycin (50 μg/ml) at 37 °C until an OD_600_ between 0.6 and 0.8 was reached. Protein overexpression was induced ON at 30 °C with 0.2 mM IPTG (US Biological, USA) and agitation at 250 rpm. Cells were harvested by centrifugation (4,500 g, 10 min, 4°C) and stored at −20°C.

Cells were resuspended in buffer A (50 mM Tris–HCl pH 8.0, 400 mM NaCl, 5% glycerol, 0.1 mM PMSF) supplemented with 20 mM imidazole and 2 mM β-mercaptoethanol. Sonication was performed on ice on a QSonica Q125 sonicator (10 s ON, 20 s OFF for a total of 15 min at 40% power) and then centrifuged (12,500 g, 30 min, 4°C). The supernatant was loaded onto a pre-equilibrated 1mL HisTrap HP column (Cytiva, USA) with buffer A. The column was washed with 10 mL of buffer A supplemented with 20 mM imidazole. The fractions were eluted with a linear gradient of buffer A supplemented with 250 mM imidazole. After gel analysis, the fraction with the purified proteins were collected, and stored at −80°C with 40% glycerol.

### Circular dichroism

The same day of circular dichroism (CD) spectra collection, the proteins were buffer-exchanged using a PD MiniTrap G-25 desalting column (Cytiva, USA) with a buffer containing 20 mM sodium phosphate pH 7.8 and Na_2_SO_4_. The fraction recovered was concentrated until reaching the desired concentration for the assay (∼13 μM). The collection of CD spectra was done on a JASCO J-1500 (Jasco Inc., Japan) at 20 nm/min with a step size of 1 nm in a 1 mm quartz cuvette using 200 μL of sample. Three scans were made per sample. The baseline of the buffer was subtracted to the average spectra of the samples. Units of Mean Residue Molar Ellipticity ([θ]*_MR_*) were calculated according to the following equation:

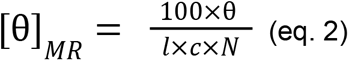

where *l* is the cell pathlength in cm, *c* is the molar concentration in M, and *N* is the number of amino acids.

A theoretical CD spectra for the best-ranked AF2-predicted structure of *P. atlantica* RfaH was calculated using SESCA v0.97 (Nagy *et al*, 2019). The resulting theoretical spectra of *P. atlantica* RfaH, together with the experimental CD spectra and the deposited PDB structures of *E. coli RfaH* and NusG (RfaH: PDB ID 5OND, chain A; NusG, PDB ID 6GOV, chain G), were processed with the BeStSel webserver (https://bestsel.elte.hu/index.php) (Micsonai *et al*, 2025) to obtain the percent of helix (regular helix + distorted helix) and strand (parallel + antiparallel) content for each sample, which was used to calculate the ratio of helix/strand (%α/%β) for each protein.

## DATA AVAILABILITY

The datasets and computer code produced in this study are available in the following database: - Protein structure predictions of RfaH orthologs using ColabFold, TMD decoys for AF2Rank, Jupyter Notebooks for reproducing our data analysis using Google Colab and results from the analysis of all protein structures: Zenodo (https://doi.org/10.5281/zenodo.19041373).

Plasmids generated in this work are available from the corresponding authors upon request.

## ACKNOWLEDGMENT

This research was funded by the National Agency for Research and Development (ANID) through Fondo Nacional de Desarrollo Científico y Tecnológico (FONDECYT Regular 1240205 to CARS, FONDECYT Postdoctorado 3240319 to PGD, FONDECYT Postdoctorado 3250787 to VDP, ANID PFCHA 21231023 to CTA, ANID PFCHA 21191684 to JR, ANID PFCHA 21230688 to CNM), the ANID Millennium Science Initiative Program (ICN17_022 to CARS), and the National Institutes of Health (NIH) grant R01 GM067153 to IA. The CD spectra were measured using a Jasco J-1500 spectropolarimeter funded by ANID FONDEQUIP EQM140151 and available at Centro de Estudios para el Desarrollo de la Química (CEPEDEQ) from Universidad de Chile. Powered@NLHPC: This research was partially supported by the supercomputing infrastructure of the NLHPC (CCSS210001). We thank the assistance of Prof. Exequiel Medina from Universidad de Chile for his help in performing the CD measurements.

## DISCLOSURE AND COMPETING INTEREST STATEMENT

C.A.R-S. is co-founder of and scientific advisor to NexEnzymes and owns equity in NexEnzymes SpA. All other authors declare no competing interests.

## Notes

https://dx.doi.org/10.5281/zenodo.19041374

### Summary of Updates

Figure 5 revised due to some typos in the p-value lettering; new results in Figure 7 of circular dichroism experiments from putative monomorphic RfaH orthologs; changing previous Figure 7 to Figure 8; new methods describing overexpression, solubility test and purification of monomorphic RfaH orthologs and circular dichroism experiments; new disclosure and competing interest statement; supplemental files updated

