## Supplementary Information Appendix for "Exploration of the structural and functional diversity in the metamorphic RfaH subfamily"

### TABLE OF CONTENTS


Appendix Figure S1 Page 2

Appendix Figure S2 Page 3

Appendix Figure S3 Page 4

Appendix Figure S4 Page 5

Appendix Figure S5 Page 6

Appendix Figure S6 Page 7

Appendix Figure S7 Page 8

Appendix Figure S8 Pages 9-10

Appendix Figure S9 Page 11

Appendix Figure S10 Page 12

Appendix FIgure S11 Page 13

Appendix Figure S12 Page 14

Appendix Figure S13 Page 15

Appendix Figure S14 Page 16

Appendix Figure S15 Page 17

Appendix Figure S16 Page 18

Appendix Table S1 Page 19

Appendix Table S2 Pages 20-21

Appendix Table S3 Pages 22-23

Appendix Table S4 Pages 24-25

Appendix Table S5 Pages 26-27

Appendix Table S6 Pages 28-29

Appendix Table S7 Pages 30-31

Appendix Table S8 Page 32

Appendix References Page 33

### APPENDIX FIGURES

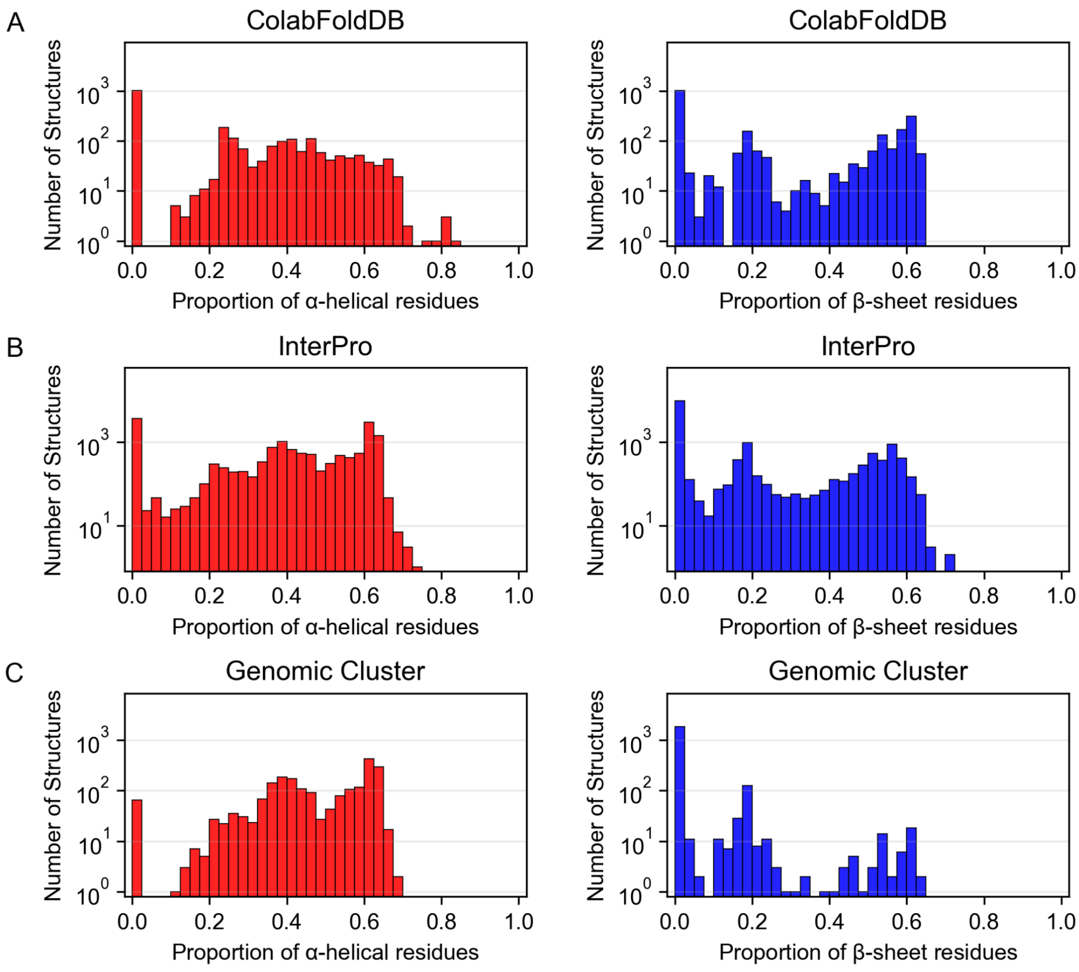


**Appendix Figure S1** Distribution of secondary structure in the CTD of structure predictions of full-length RfaH homologs. Distribution of α-helical (red) and β-strand (blue) secondary structure content analyzed using STRIDE for the 19,710 predicted structures of ColabFoldDB (**A**, 2,340 structures), InterPro (**B**, 15,290 structures), and Genomic Cluster (**C**, 2,080 structures), with the y axis corresponding to the number of structures with a given proportion of secondary structure in logarithmic scale. A small dip in the proportion of secondary structure is observed at 0.325 for α-helical residues and 0.300 for β-strands (more clearly observed for the plots in **C**). We used these fractions as thresholds to distinguish between αRfaH (fraction of α-helical structure > 0.325 and β-strand ~0), βRfaH (fraction of β-strand structure > 0.300 and α-helical ~0) or mixed α/β secondary structure (fraction of α-helical structure and β-strand structure > 0.025).

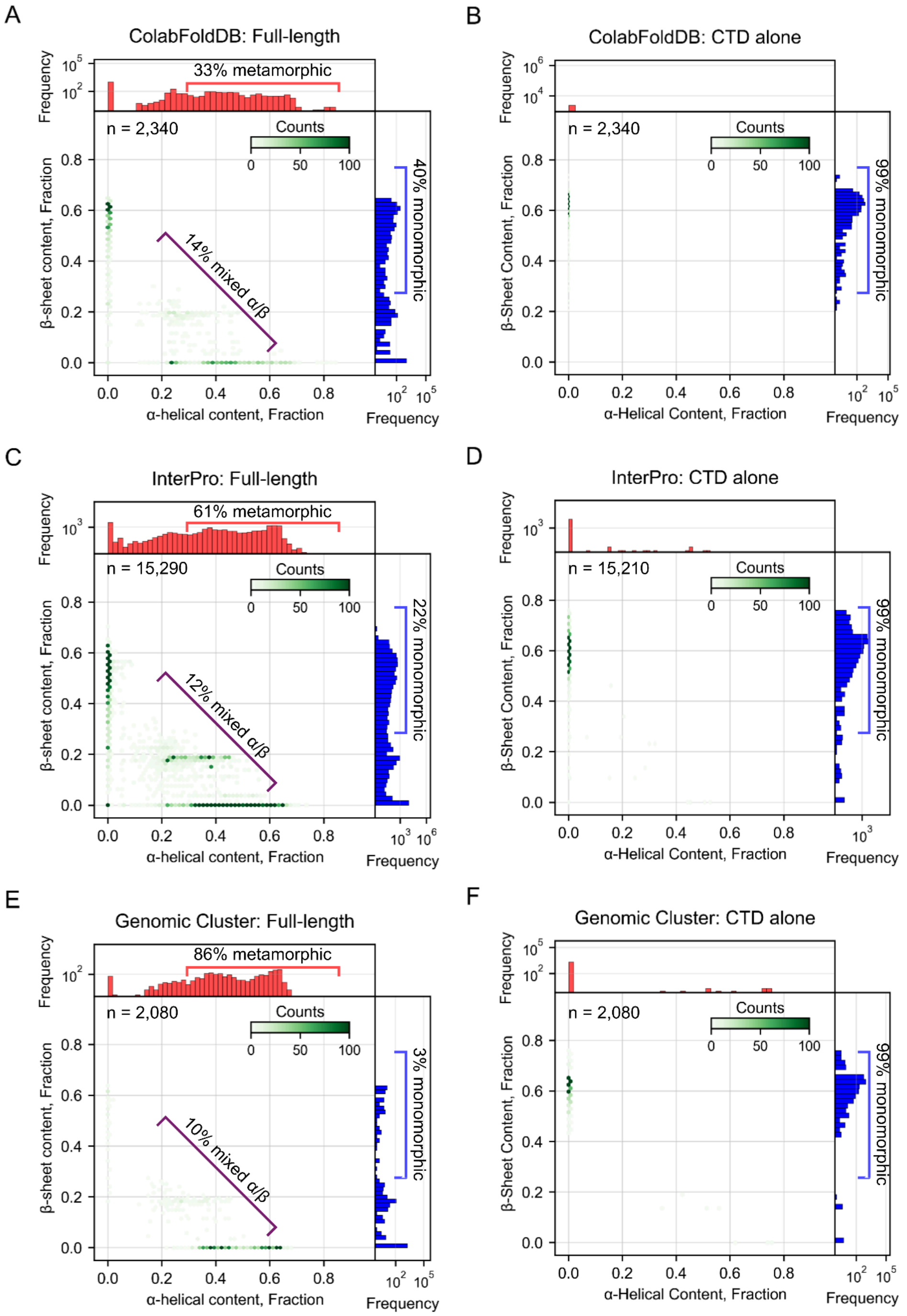


**Appendix Figure S2** Identification of metamorphic, mixed α/β CTD, and monomorphic RfaH homologs based on AF2 protein structure predictions. Secondary structure content of predictions of RfaH homologs from ColabFoldDB (**A**, **B**), InterPro (**C**, **D**), and a reported Genomic Cluster (**E**, **F**). Predictions were made using either the full-length RfaH sequences from each database (**A**, **C**, **E**) or the isolated CTD (**B**, **D**, **F**), determined based on MSAs generated using MAFFT. The secondary structure assignment of the CTD, determined using STRIDE, shows groups of predicted RfaH structures that display an αCTD, a βCTD, or mixed α/β secondary structure. Some structures from InterPro had short CTD sequences.

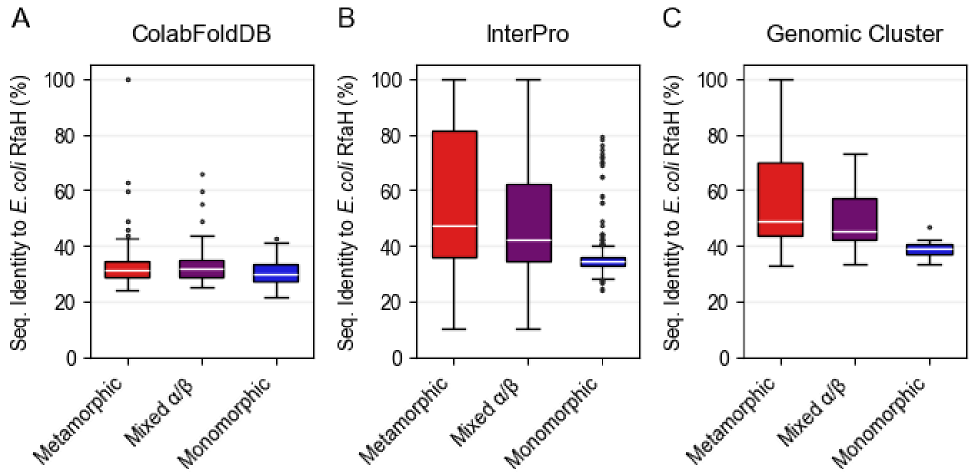


**Appendix Figure S3** Distribution of sequence identity against *E. coli* RfaH for all datasets. The structural predictions from the ColabFoldDB (**A**), InterPro (**B**) and (**C**) Genomic cluster databases presented in Figure 2 were used to assign each structure to a CTD fold category, namely αCTD (red), βCTD (blue), or mixed α/β CTD (purple), and its sequence identity calculated against *E. coli* RfaH was appended to the corresponding category. Each sequence can be present at most once in every category, based on its predicted CTD. In the box plots, the median is shown as a horizontal white line, the lower and upper ends indicate the 25% and 75% interquartile range, and the whiskers extend to the percentile 1 and 99, with points representing outliers.

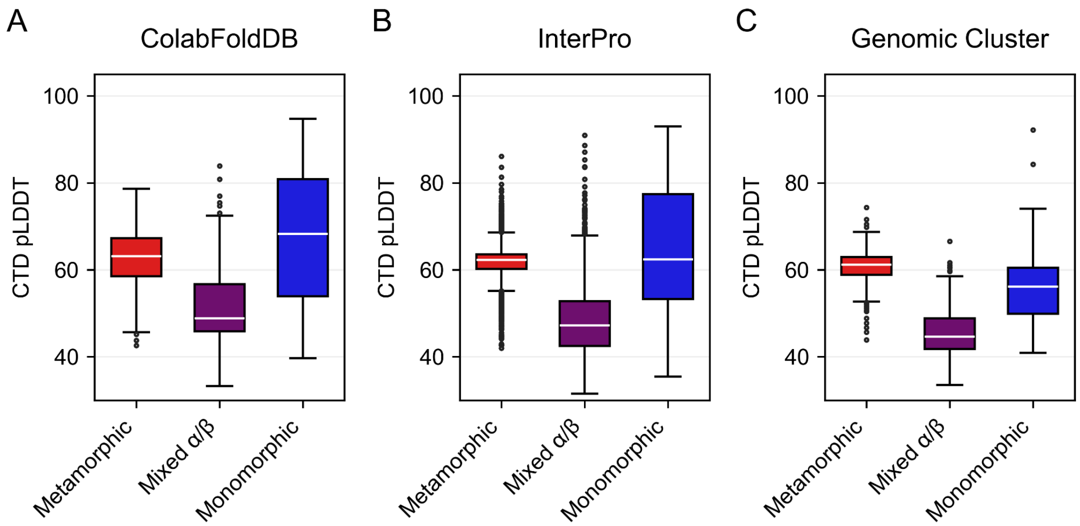


**Appendix Figure S4** Distribution of the pLDDT of the CTD of full-length RfaH protein structures from all databases. The average pLDDT of the CTD from the RfaH structures predicted form the ColabFoldDB (**A**), InterPro (**B**) and Genomic Cluster (**C**) datasets was used as a proxy to rank the best predicted structures, and these values are presented in these plots clustered by CTD fold category, namely αCTD (red), βCTD (blue), or mixed α/β CTD (purple). Each sequence can be present at most once in every category, based on its predicted CTD. In the box plots, the median is shown as a horizontal white line, the lower and upper ends indicate the 25% and 75% interquartile range, and the whiskers extend to the percentile 1 and 99, with points representing outliers.

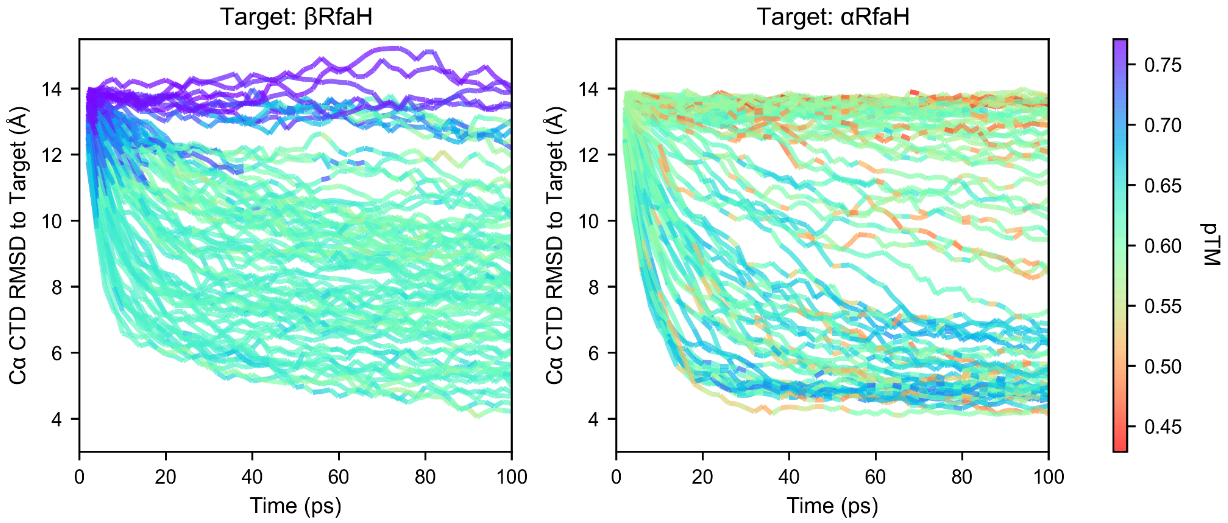


**Appendix Figure S5** Graphical representation of the TMD trajectories. A total of 200 TMD trajectories were produced, in which backbone atoms from residues 100-162, comprising *E. coli* RfaH CTD were subjected to an external potential to fold-switch from the autoinhibited (**A**) or the active state (**B**) towards the target opposite native state, and ~100 simulations were selected based on Principal Component Analysis to create thousands of decoys for further analysis using AF2Rank. Each line represents a single trajectory of 100 ps, with the colors indicating the predicted TM score (pTM) for each input structure. The plots show the Cα RMSD of *E. coli* RfaH CTD against the target structure, indicated in the title of each plot.

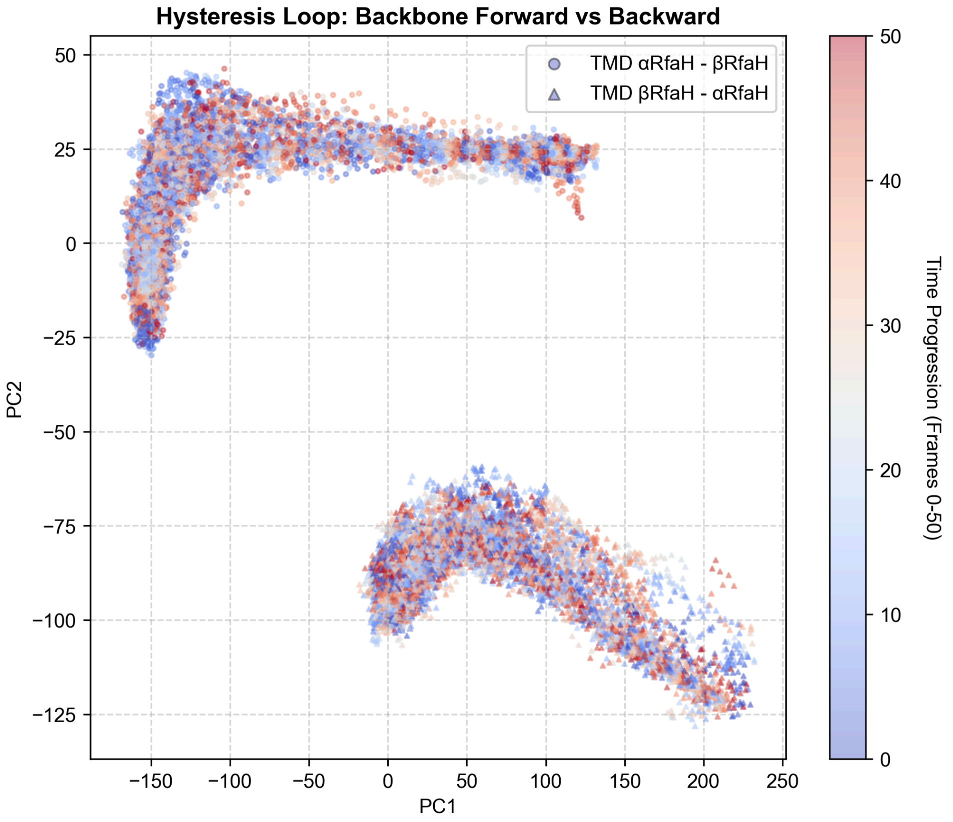


**Appendix Figure S6** Hysteresis of the TMD simulations. Principal Component Analysis (PCA) projections based on a coordinate covariance matrix, which was calculated using the Cα atoms of the CTD (residues 110–162) of *E. coli* RfaH, while the stable NTD (residues 1–100) was used as a rigid-body reference for alignment to remove translational and rotational motions. The covariance matrix was diagonalized to obtain the eigenvectors and project all 600 TMD trajectories (300 in each fold-switching direction) based on two components, PC1 and PC2, and the trajectory frames were clustered into at least 100 distinct groups based on their Euclidean distance in the PC1/PC2 subspace using *k*-means clustering. The centroid of each cluster was identified, and the specific trajectory containing the centroid frame was selected as a representative pathway. This process yielded a curated dataset of 100 simulations for the TMD trajectories from αRfaH to βRfaH and 99 trajectories from βRfaH to αRfaH.

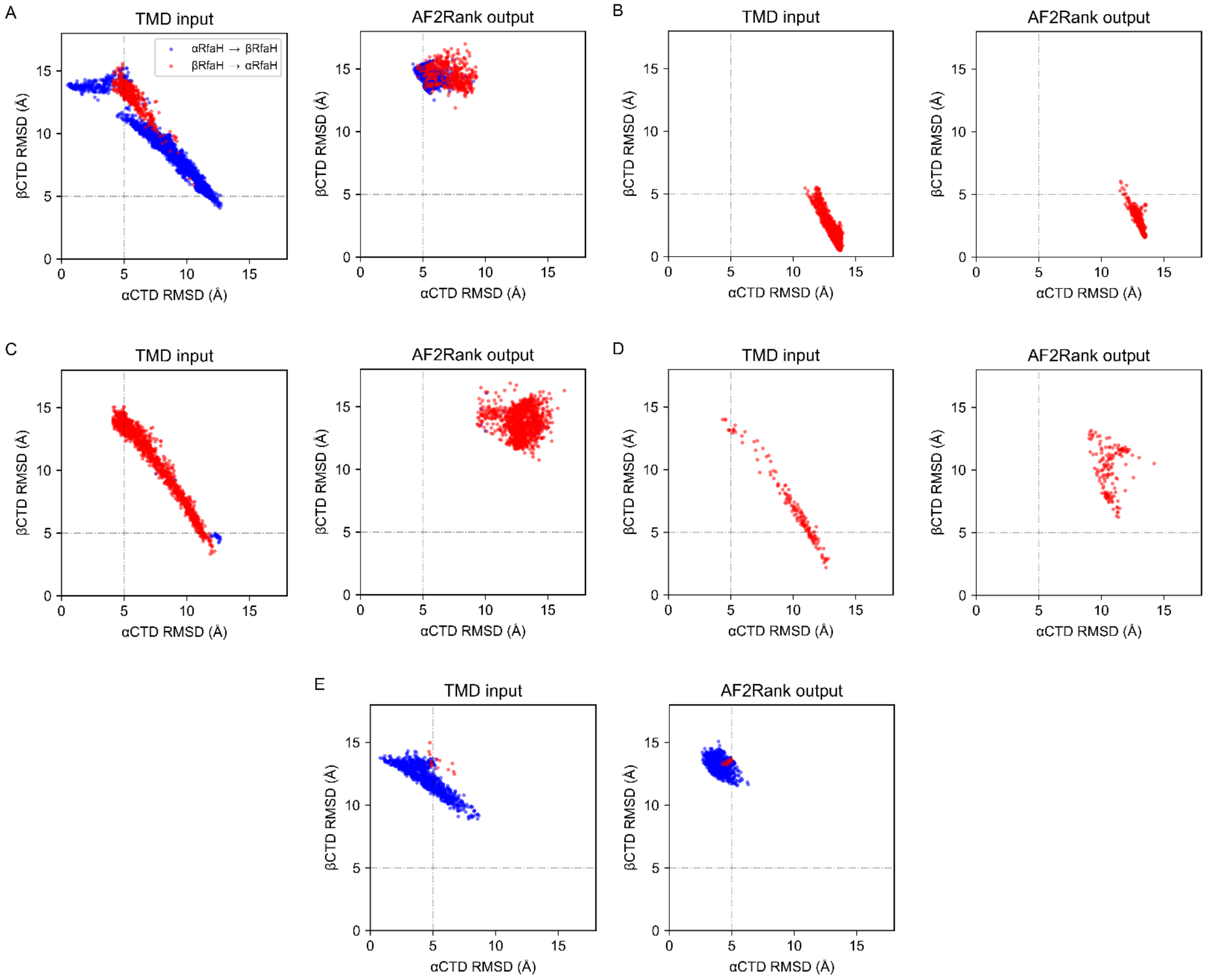


**Appendix Figure S7** Structural mapping of the input decoys generated by TMD and the output structures refined by AF2Rank. Cluster 1 (**A**) contains AF2Rank output structures generated from 2,471 decoys from TMD simulations from αRfaH to βRfaH and 472 decoys from the reverse simulations; cluster 2 (**B**) contains AF2Rank output structures from 1,704 input structures from TMD simulations from βRfaH to αRfaH; cluster 3 (**C**) contains AF2Rank output structures generated from 23 decoys from TMD simulations from αRfaH to βRfaH and 1,357 decoys from the reverse simulations; cluster 4 (**D**) contains AF2Rank output structures from 153 input structures from TMD simulations from βRfaH to αRfaH; and cluster 5 (**E**) contains AF2Rank output structures generated from 986 decoys from TMD simulations from αRfaH to βRfaH and 14 decoys from the reverse simulations. Clusters 1, 3 and 4 represent intermediate states with inverted helices (cluster 1), and mixed secondary structure (groups 3 and 4), as shown in Figure 4 in the main text, whereas clusters 2 and 5 resemble the active and autoinhibited states of *E. coli* RfaH, respectively.

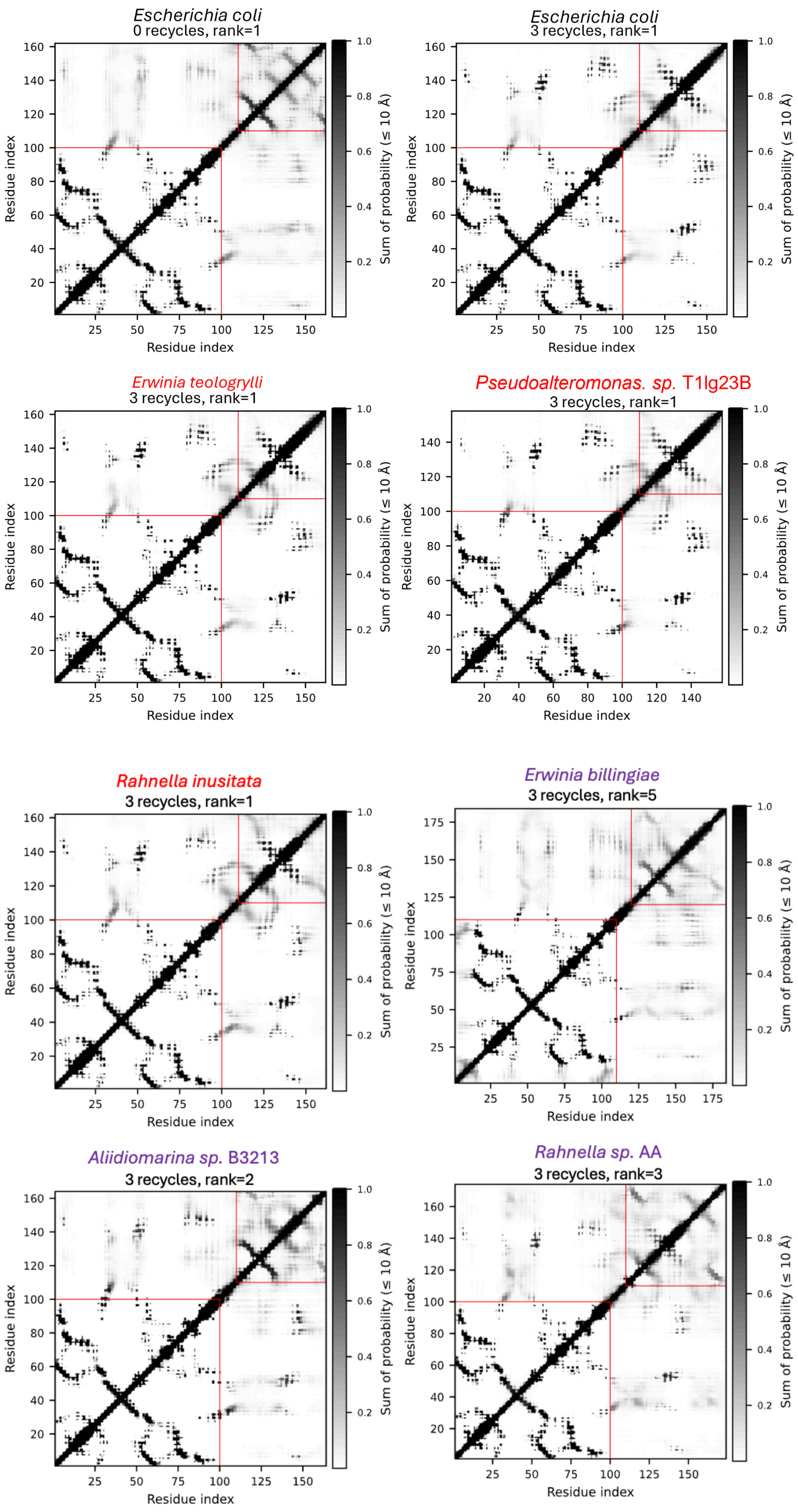

Escherichia coli
0 recycles, rank=1
Escherichia coli
3 recycles, rank=1
Erwinia teologrylli
3 recycles, rank=1
Pseudoalteromonas. sp. T1lg23B
3 recycles, rank=1
Rahnella inusitata
3 recycles, rank=1
Erwinia billingiae
3 recycles, rank=5
Aliidiomarina sp. B3213
3 recycles, rank=2
Rahnella sp. AA
3 recycles, rank=3
Residue index
Sum of probability (≤ 10 Å)

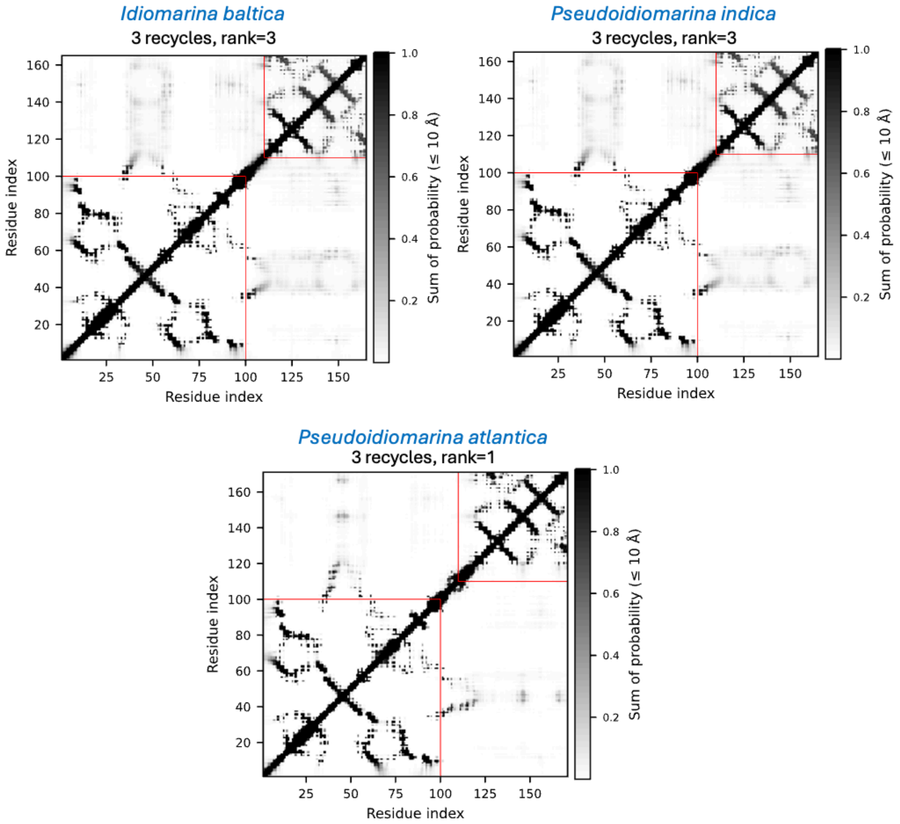


**Appendix Figure S8** Distograms for *E. coli* RfaH in the autoinhibited and active states and metamorphic, mixed α/β and monomorphic RfaH orthologs AF2 predictions from the Genomic Cluster. Distograms plot the contact probability up to 10 Å cutoff for RfaH orthologs predicted by AF2, highlighting the species name in the color code: *E. coli* (best ranked structures from ColabFold predictions at recycle 0 and recycle 3) in black, metamorphic orthologs in red, RfaH orthologs with mixed α/β CTD in purple, and RfaH homologs predicted as monomorphic in the canonical NusG-like active state in blue.

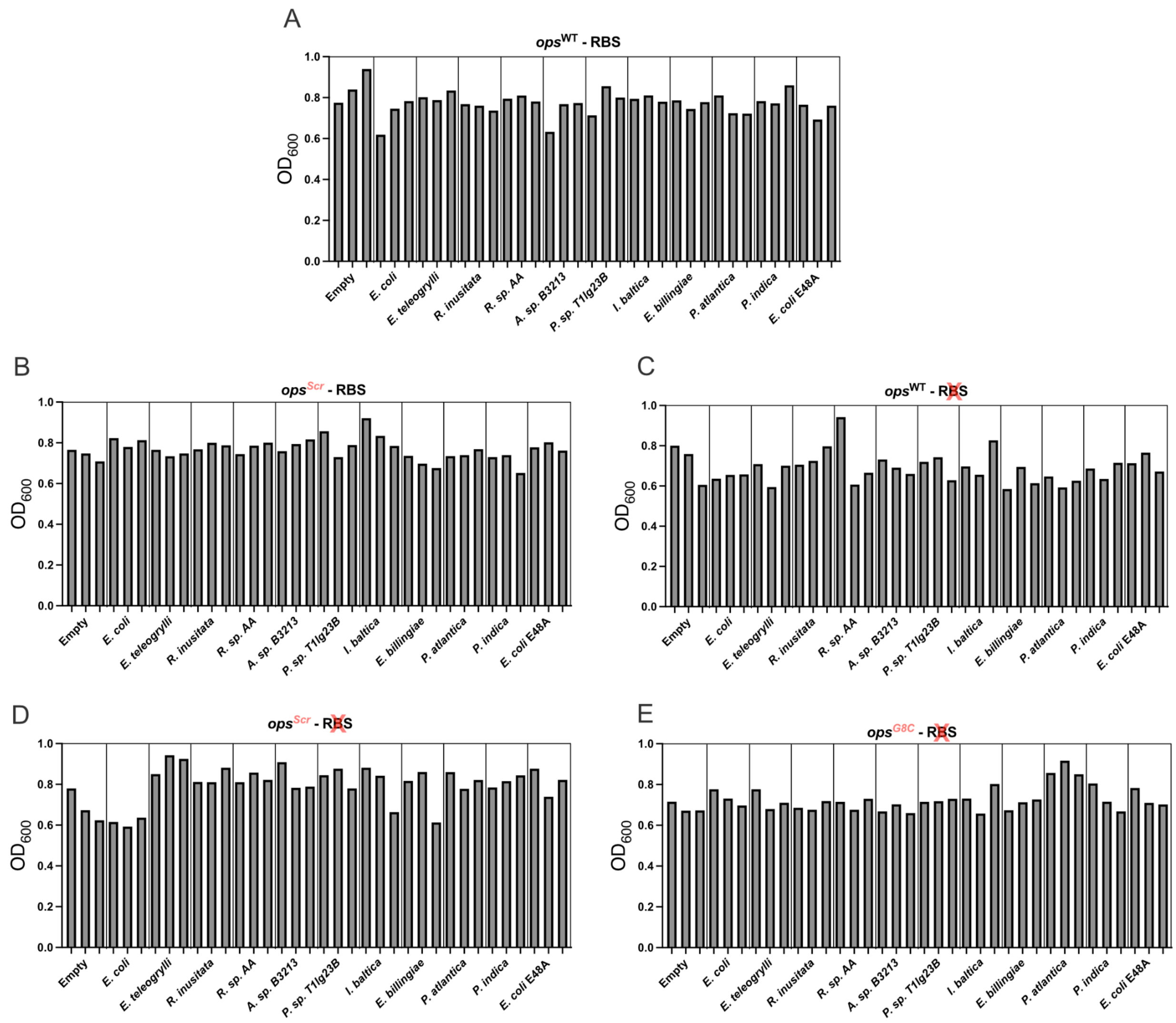


**Appendix Figure S9** Final $OD_{600}$ measured for each colony used in the *in vivo* luminescence assays under non-inducible conditions. The genetic architectures for the plasmid encoding the luminescence reporter correspond to *ops*$^{WT}$ with RBS (**A**), *ops*$^{Scr}$ with RBS (**B**), *ops*$^{WT}$ without RBS (**C**), *ops*$^{Scr}$ without RBS (**D**) and *ops*$^{G8C}$ without RBS (**D**). Three biological replicates were performed per RfaH protein per condition.

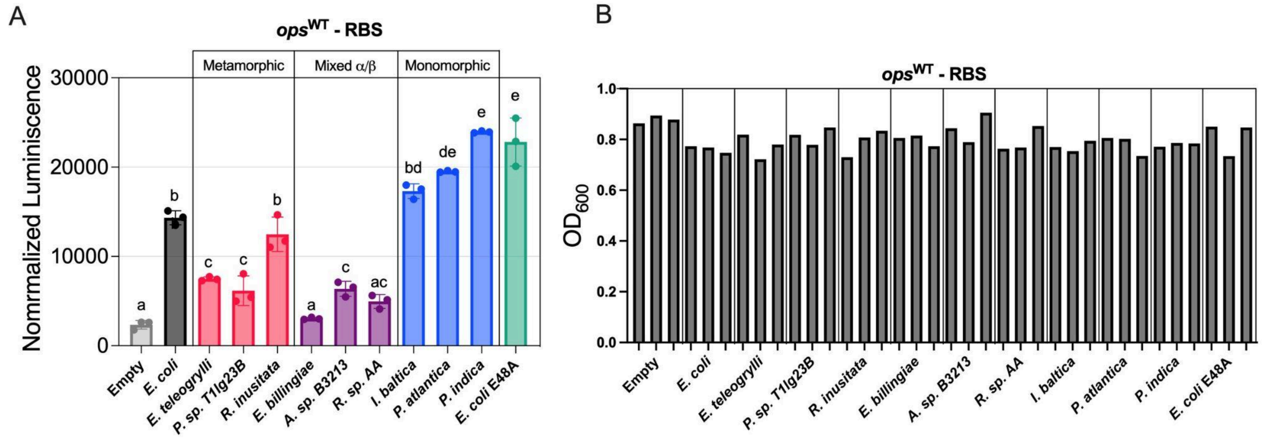


**Appendix Figure S10** *In vivo lux* assay under IPTG induction of expression of RfaH homologs. (**A**) The plasmid encoding the *luxCDABE* operon downstream of the wild-type *ops*-element and RBS was co-transformed with the plasmid encoding the RfaH-like proteins in *E. coli* DH5α Δ*rfaH* cells or with an empty vector as control. The *in vivo* luminescence assay was carried out using IPTG concentrations to induce RfaH overexpression (see *Methods*). Each bar represents the average raw luminescence level of six technical replicates of three colonies, normalised by cell density. The error bar shows the standard deviation. A one-way Analysis of Variance (ANOVA) was used to assess statistical significance, with all samples compared to each other. The same letter means non-significant different (p-value > 0.05), while different letter means statistical difference (p-value ≤ 0.05). (**B**) Final $OD_{600}$ measured for each colony used in the assay. The raw luminescence of each colony was normalized to its corresponding final $OD^{600}$. Three biological replicates were performed per RfaH protein per condition.

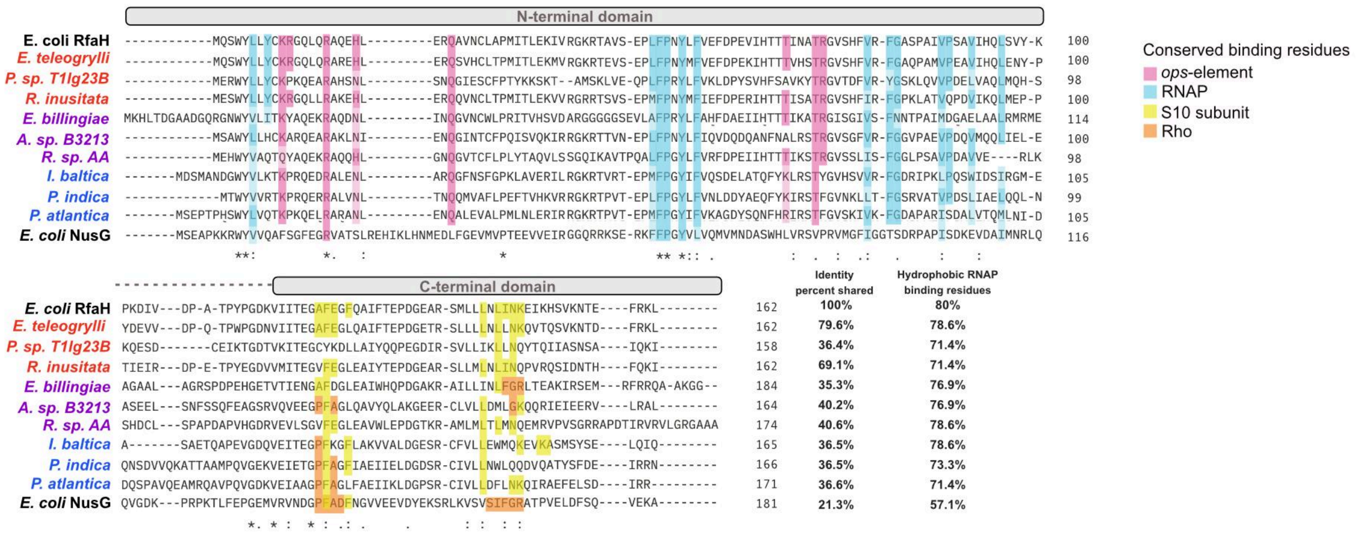


**Appendix Figure S11** MSA of RfaH orthologs tested in the *lux* assay. The MSA was generated using MUSCLE (Edgar, 2004) available at the EMBL-EBI Job Dispatcher web server (Madeira *et al*, 2024). The orthologs classified as metamorphic, mixed α/β CTD and monomorphic are labeled in red, purple, and blue, respectively. The MSA was built including the sequences of *E. coli* RfaH and NusG as references. The colors in the columns indicate the conservation of residues associated with binding to the *ops* DNA element (pink), RNAP (cyan), the S10 subunit (yellow) and Rho (orange). It can be observed that there is only partial conservation of *ops*-binding residues and of residues from loop 1 involved in Rho binding in the case of the monomorphic proteins.

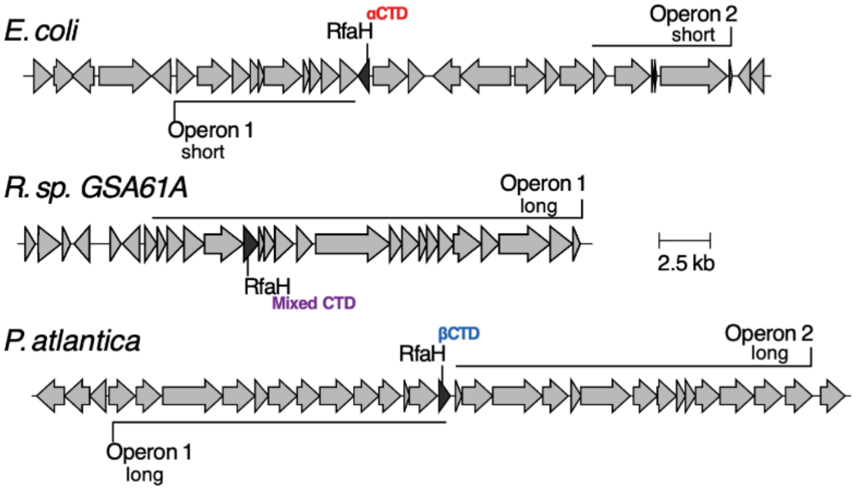


**Appendix Figure S12** Gene neighborhood of genomic-encoded RfaH orthologs. Representative genomic vicinity of the metamorphic, mixed α/β CTD and monomorphic RfaH experimentally ascertained in this work. The surrounding 15 genes up and downstream of the *rfaH* gene are shown, as well as their open reading frame (ORF) direction, for three organisms: *E. coli* RfaH, the canonical metamorphic protein; *Rhanella sp.* RfaH, belonging to the mixed α/β CTD clade; and *P. atlantica* RfaH, belonging to the monomorphic clade. Based on the operon definition described in the *Methods* section, different operon lengths near or away from the RfaH production site can be observed among these three examples, with the mixed α/β CTD of *Rhanella sp.* and the monomorphic RfaH from *P. atlantica* being part of an operon and acting in *cis*, whereas *E. coli* contains operons for which RfaH must act in *trans*.

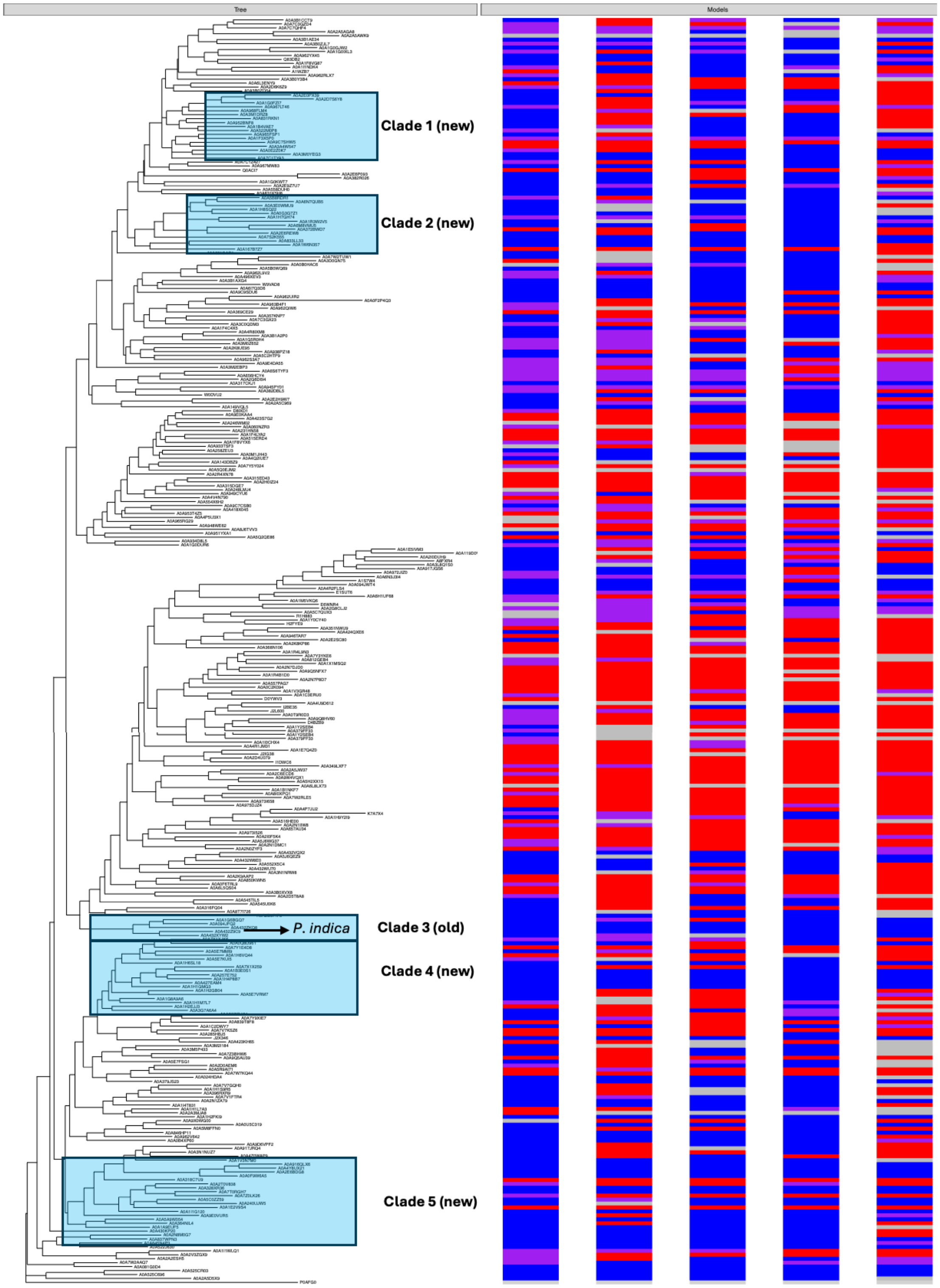


**Appendix Figure S13** Genbank-matched InterPro sequences and dendrogram for distant RfaH orthologs. In this dendrogram, 5 groups of sequences (Clade 1-5) are characterized as putative monomorphic RfaH homologs. The results of their genomic context are summarized in Appendix Table 3.

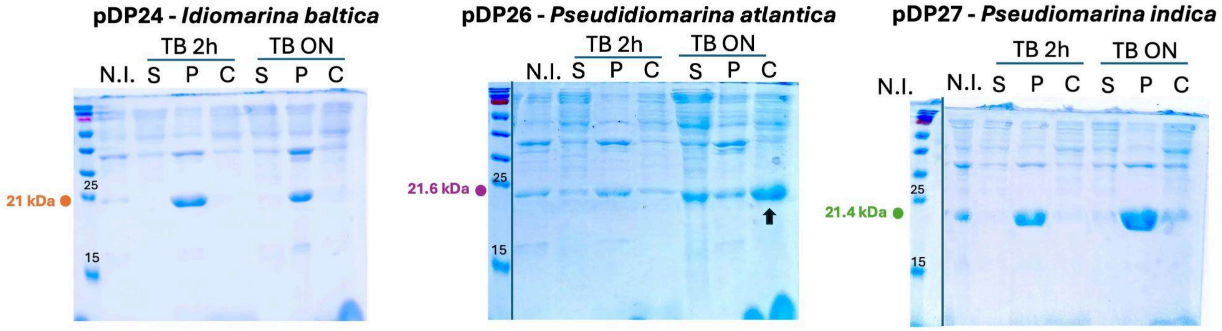


**Appendix Figure S14** Overexpression and solubility test performed on RfaH orthologs predicted as putative monomorphic proteins. The cells were grown in TB medium supplemented with kanamycin. After reaching the desired $OD_{600}$, samples were induced for a short (2 h) and long period of time (ON). The samples obtained in were run in a 12% SDS-PAGE gel, for the control with no induction (N.I.), soluble fraction (S), insoluble fraction or pellet (P) and the insoluble fraction resuspended with buffer LysT 6.9 supplemented with 1% CHAPS (C). Only the putative monomorphic RfaH from *P. atlantica* showed soluble expression, indicated with a black arrow

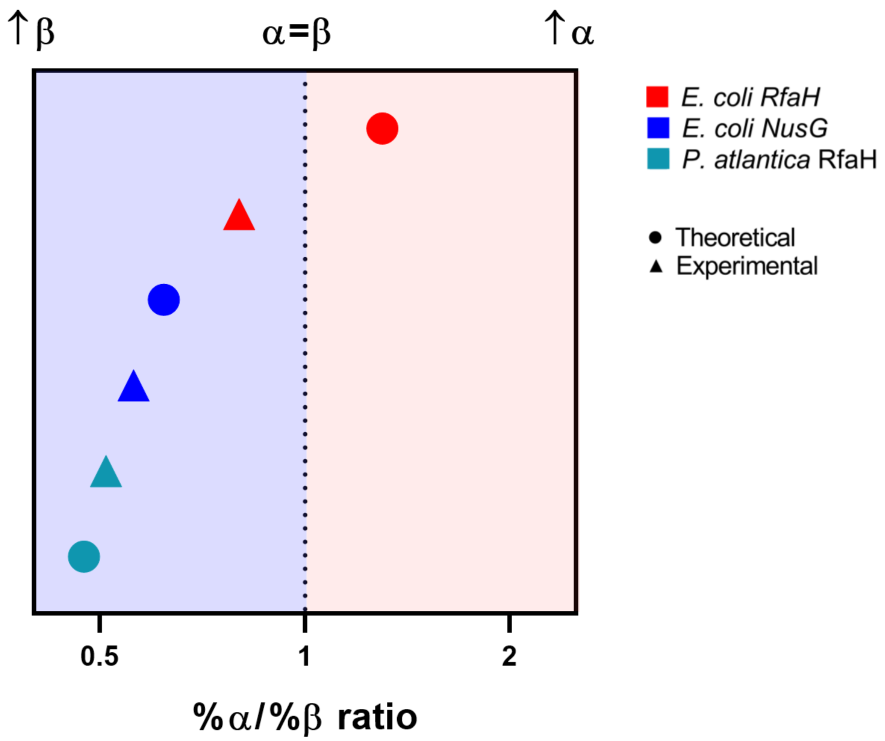


**Appendix Figure S15** Comparison of the %α/%β ratio of *P. atlantica* RfaH (teal), *E. coli* NusG (blue), and *E. coli* RfaH (red) obtained from experimental CD spectrum (▲) and theoretical data (●, PDB ID 6GOV chain G for *E. coli* NusG; PDB ID 5OND chain A for *E. coli* RfaH, and the top-ranked AF2 model for *P. atlantica* RfaH). The vertical dotted line marks a ratio of 1; to the left (blue background) β-sheet structure predominates (↑β), and to the right (red background) α-helix structure predominates (↑α). The relative ordering of the proteins is consistent across all three data types. The %α/%β ratio was obtained with the BeStSel server: directly from the empirical CD spectrum (▲), or from the crystal structure or a theoretical CD spectrum generated with SESCA v0.97 from the predicted structure (●).

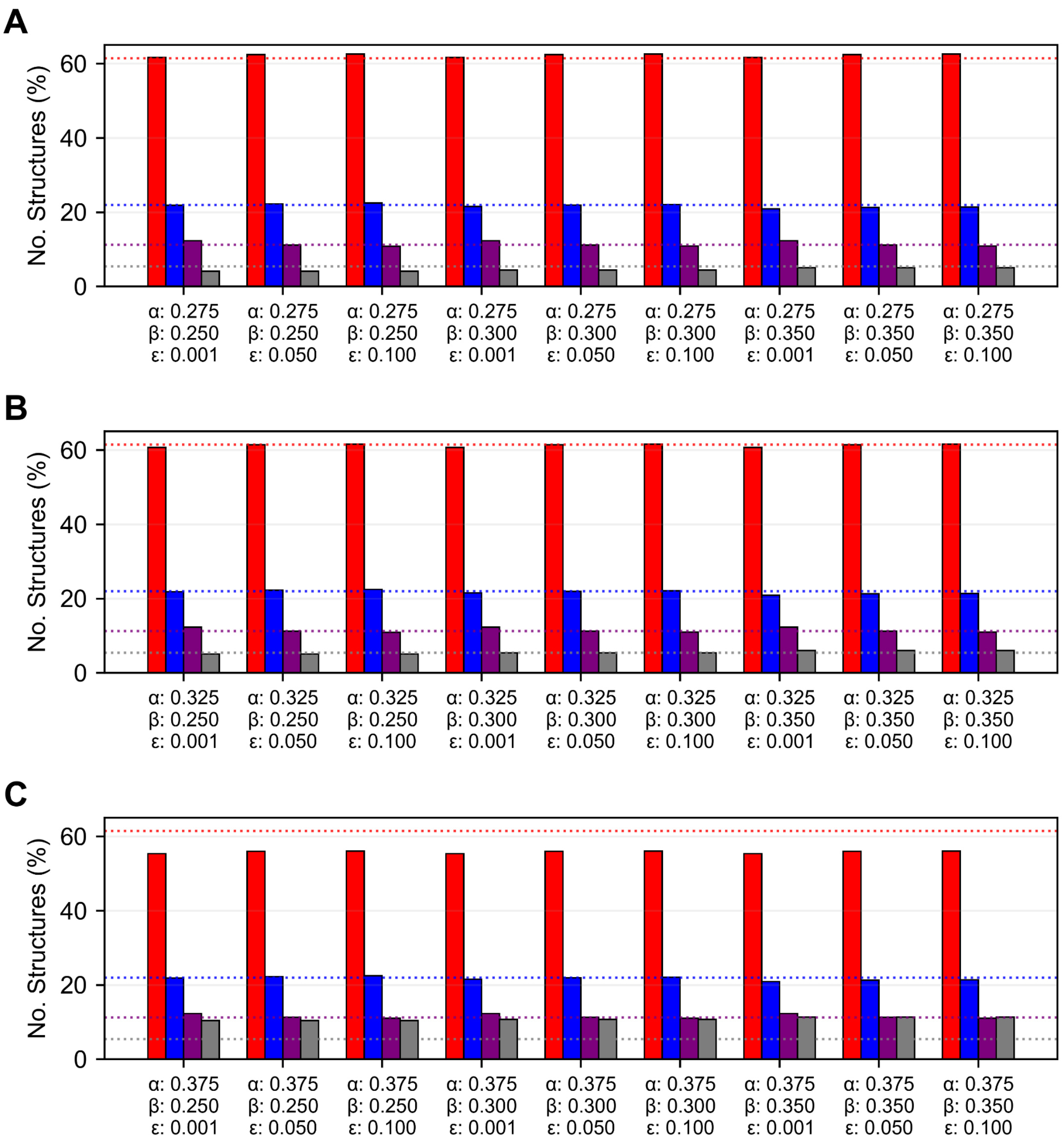


**Appendix Figure S16** Sensitivity test of the thresholds used to distinguish between metamorphic, monomorphic and mixed α/β CTDs in full-length RfaH orthologs. The used thresholds (metamorphic αRfaH: α-helices [α] > 32.5% and < 2.5% β-strands [ε]; monomorphic βRfaH: β-strands [β] > 30.0% and < 2.5% α-helices [ε]; mixed α/β CTD with α-helices and β-strands > 2.5% [ε]) which lead to the number of structures indicated by dotted lines for metamorphic (red), monomorphic (blue), mixed (purple) and uncategorized structures (gray) were changed by ±5% and the variation in the number of structures per classification was determined for all structures predicted from all databases. In (**A**), only α was fixed to 27.5%, in (**B**), only α was fixed to 32.5%, and in (**C**) only α was fixed to 37.5%. Only the metamorphic proteins show a decrease in the total number of structures upon increasing the threshold for α-helices > 37.5%, with a concomitant increase in uncategorized structures from 4 to 10%, whereas monomorphic structures (blue) and mixed α/β structures show minimal changes.

### APPENDIX TABLES

**Appendix Table S1** Summary of selected RfaH proteins for *in vivo* luminescence assay.

| Organism | GenBank ID | AF2 Prediction | Topology | Sequence identity against *E. coli* RfaH |
|---|---|---|---|---|
| *Escherichia coli* | NP_418284 | 5α 0α/β 0β | Metamorphic | 100% |
| *Erwinia teleogrylli* | WP_058911819.1 | 5α 0α/β 0β | Metamorphic | 79.6% |
| *Pseudoalteromonas. sp.* T1lg23B | WP_105167350.1 | 5α 0α/β 0β | Metamorphic | 36.4% |
| *Rahnella inusitata* | WP_112168251.1 | 3α 2α/β 0β | Metamorphic | 69.1% |
| *Erwinia billingiae* | WP_013200013.1 | 1α 2α/β 2β | Mixed α/β | 35.3% |
| *Aliidiomarina sp.* B3213 | WP_111914955.1 | 2α 2α/β 1β | Mixed α/β | 40.2% |
| *Rahnella sp.* AA | WP_101080087.1 | 1α 4α/β 0β | Mixed α/β | 40.6% |
| *Idiomarina baltica* | WP_006956899.1 | 2α 1α/β 2β | Monomorphic | 36.5% |
| *Pseudidiomarina indica* | WP_092591975.1 | 1α 0α/β 4β | Monomorphic | 36.5% |
| *Pseudidiomarina atlantica* | WP_034731719.1 | 0α 0α/β 5β | Monomorphic | 36.6% |

**Appendix Table S2** Plasmids used in this work

| Plasmids | Name | Description | Resistance | Reference |
|---|---|---|---|---|
| *ops* and RBS variants for *lux* assay | pIA955 | $P_{BAD}$–*ops*$^{WT}$_RBS-*luxCDABE* | Amp | (Belogurov *et al*, 2010) |
| | pIA1087 | $P_{BAD}$–*ops*$^{WT}$–*luxCDABE* | Amp | (Burmann *et al*, 2012) |
| | pZL23 | $P_{BAD}$–*ops*$^{G8C}$–*luxCDABE* | Amp | (Zuber *et al*, 2018) |
| | pZL4 | $P_{BAD}$–*ops*$^{Scr}$_RBS-*luxCDABE* | Amp | (Zuber *et al*, 2018) |
| | pZL8 | $P_{BAD}$–*ops*$^{Scr}$–*luxCDABE* | Amp | (Zuber *et al*, 2018) |
| Expression vector for *lux* assay | pIA947 | $P_{trc}$- no CDS ("Empty") | Cm | (Belogurov *et al*, 2010) |
| | pIA957 | $P_{trc}$- RfaH$^{E. coli}$ | Cm | (Belogurov *et al*, 2010) |
| | pIA1072 | $P_{trc}$- RfaH$^{E. coli}$ (E48A) | Cm | (Burmann *et al*, 2012) |
| | pDP10 | $P_{trc}$- RfaH$^{E. teologrylli}$ | Cm | This work |
| | pDP11 | $P_{trc}$- RfaH$^{R. inusitata}$ | Cm | This work |
| | pDP12 | $P_{trc}$- RfaH$^{R. sp. AA}$ | Cm | This work |
| | pDP13 | $P_{trc}$- RfaH$^{A. sp. B3213}$ | Cm | This work |
| | pDP14 | $P_{trc}$- RfaH$^{P. sp. T1lg23B}$ | Cm | This work |
| | pDP15 | $P_{trc}$- RfaH$^{Idiomarina\ baltica}$ | Cm | This work |
| | pDP16 | $P_{trc}$- RfaH$^{E. billingiae}$ | Cm | This work |
| | pDP17 | $P_{trc}$- RfaH$^{P. atlantica}$ | Cm | This work |
| | pDP18 | $P_{trc}$- RfaH$^{P. indica}$ | Cm | This work |
| Overexpression vector for | pIA244 | $P_{T7}$-NusG$^{E. coli}$-His$_6$ | Kan | (Belogurov *et al*, 2007) |

| | | | | |
|---|---|---|---|---|
| protein purification | EcoRfaH | $P_{T7}$- $His_6$-$RfaH^{E. coli}$ | Kan | This work |
| | pDP24 | $P_{T7}$- $His_6$-$RfaH^{I. baltica}$ | Kan | This work |
| | pDP26 | $P_{T7}$- $His_6$-$RfaH^{P. atlantica}$ | Kan | This work |
| | pDP27 | $P_{T7}$- $His_6$-$RfaH^{P. indica}$ | Kan | This work |

**Appendix Table S3** p-values from one-way ANOVA Multiple Comparison of Figure 5B

| Comparison | Adjusted p-value | Summary | Letter group 1 | Letter group 2 |
|---|---|---|---|---|
| Empty vs. *E. coli* | 0.0003 | *** | a | bc |
| Empty vs. *E. teleogrylli* | >0.9999 | ns | a | a |
| Empty *vs. R. inusitata* | 0.3442 | ns | a | ab |
| Empty *vs. R. sp. AA* | >0.9999 | ns | a | a |
| Empty *vs. A. sp. B3213* | 0.6704 | ns | a | a |
| Empty *vs. P. sp. T1lg23B* | 0.9592 | ns | a | a |
| Empty *vs. I. baltica* | 0.0001 | *** | a | bc |
| Empty *vs. E. billingiae* | >0.9999 | ns | a | a |
| Empty *vs. P. atlantica* | <0.0001 | **** | a | c |
| Empty *vs. P. indica* | <0.0001 | **** | a | d |
| Empty *vs. E. coli E48A* | <0.0001 | **** | a | d |
| *E. coli vs. E. teleogrylli* | 0.0003 | *** | bc | a |
| *E. coli vs. R. inusitata* | 0.1209 | ns | bc | ab |
| *E. coli vs. R. sp. AA* | 0.0004 | *** | bc | a |
| *E. coli vs. A. sp. B3213* | 0.0388 | * | bc | a |
| *E. coli vs. P. sp. T1lg23B* | 0.0085 | ** | bc | a |
| *E. coli vs. I. baltica* | >0.9999 | ns | bc | bc |
| *E. coli vs. E. billingiae* | <0.0001 | **** | bc | a |
| *E. coli vs. P. atlantica* | 0.6816 | ns | bc | c |
| *E. coli vs. P. indica* | <0.0001 | **** | bc | d |
| *E. coli vs. E. coli E48A* | <0.0001 | **** | bc | d |
| *E. teleogrylli vs. R. inusitata* | 0.2969 | ns | a | ab |
| *E. teleogrylli vs. R. sp. AA* | >0.9999 | ns | a | a |
| *E. teleogrylli vs. A. sp. B3213* | 0.6106 | ns | a | a |
| *E. teleogrylli vs. P. sp. T1lg23B* | 0.9368 | ns | a | a |
| *E. teleogrylli vs. I. baltica* | 0.0001 | *** | a | bc |
| *E. teleogrylli vs. E. billingiae* | >0.9999 | ns | a | a |
| *E. teleogrylli vs. P. atlantica* | <0.0001 | **** | a | c |
| *E. teleogrylli vs. P. indica* | <0.0001 | **** | a | d |
| *E. teleogrylli vs. E. coli E48A* | <0.0001 | **** | a | d |
| *R. inusitata vs. R. sp. AA* | 0.4141 | ns | ab | a |
| *R. inusitata vs. A. sp. B3213* | >0.9999 | ns | ab | a |
| *R. inusitata vs. P. sp. T1lg23B* | 0.9830 | ns | ab | a |
| *R. inusitata vs. I. baltica* | 0.0615 | ns | ab | bc |
| *R. inusitata vs. E. billingiae* | 0.1116 | ns | ab | a |
| *R. inusitata vs. P. atlantica* | 0.0012 | ** | ab | c |
| *R. inusitata vs. P. indica* | <0.0001 | **** | ab | d |
| *R. inusitata vs. E. coli E48A* | <0.0001 | **** | ab | d |
| *R. sp. AA vs. A. sp. B3213* | 0.7468 | ns | a | a |
| *R. sp. AA vs. P. sp. T1lg23B* | 0.9795 | ns | a | a |
| *R. sp. AA vs. I. baltica* | 0.0002 | *** | a | bc |
| *R. sp. AA vs. E. billingiae* | 0.9996 | ns | a | a |
| *R. sp. AA vs. P. atlantica* | <0.0001 | **** | a | c |
| *R. sp. AA vs. P. indica* | <0.0001 | **** | a | d |
| *R. sp. AA vs. E. coli E48A* | <0.0001 | **** | a | d |
| *A. sp. B3213 vs. P. sp. T1lg23B* | >0.9999 | ns | a | a |
| *A. sp. B3213 vs. I. baltica* | 0.0184 | * | a | bc |
| *A. sp. B3213 vs. E. billingiae* | 0.2969 | ns | a | a |
| *A. sp. B3213 vs. P. atlantica* | 0.0003 | *** | a | c |
| *A. sp. B3213 vs. P. indica* | <0.0001 | **** | a | d |

| | | | | |
|---|---|---|---|---|
| *A. sp. B3213 vs. E. coli E48A* | <0.0001 | **** | a | d |
| *P. sp. T1lg23B vs. I. baltica* | 0.0039 | ** | a | bc |
| *P. sp. T1lg23B vs. E. billingiae* | 0.6839 | ns | a | a |
| *P. sp. T1lg23B vs. P. atlantica* | <0.0001 | **** | a | c |
| *P. sp. T1lg23B vs. P. indica* | <0.0001 | **** | a | d |
| *P. sp. T1lg23B vs. E. coli E48A* | <0.0001 | **** | a | d |
| *I. baltica vs. E. billingiae* | <0.0001 | **** | bc | a |
| *I. baltica vs. P. atlantica* | 0.8599 | ns | bc | c |
| *I. baltica vs. P. indica* | <0.0001 | **** | bc | d |
| *I. baltica vs. E. coli E48A* | <0.0001 | **** | bc | d |
| *E. billingiae vs. P. atlantica* | <0.0001 | **** | a | c |
| *E. billingiae vs. P. indica* | <0.0001 | **** | a | d |
| *E. billingiae vs. E. coli E48A* | <0.0001 | **** | a | d |
| *P. atlantica vs. P. indica* | 0.0052 | ** | c | d |
| *P. atlantica vs. E. coli E48A* | <0.0001 | **** | c | d |
| *P. indica vs. E. coli E48A* | 0.2307 | ns | d | d |

**Appendix Table S4** p-values from one-way ANOVA Multiple Comparison of Figure 5C

| Comparison | Adjusted p-value | Summary | Letter group 1 | Letter group 2 |
|---|---|---|---|---|
| Empty *vs. E. coli* | 0.0090 | ** | a | b |
| Empty *vs. E. teleogrylli* | 0.4589 | ns | a | ab |
| Empty *vs. R. inusitata* | 0.0787 | ns | a | ab |
| Empty *vs. R. sp. AA* | 0.7248 | ns | a | ab |
| Empty *vs. A. sp. B3213* | 0.3857 | ns | a | ab |
| Empty *vs. P. sp. T1lg23B* | 0.1416 | ns | a | ab |
| Empty *vs. I. baltica* | <0.0001 | **** | a | c |
| Empty *vs. E. billingiae* | 0.7605 | ns | a | ab |
| Empty *vs. P. atlantica* | <0.0001 | **** | a | c |
| Empty *vs. P. indica* | <0.0001 | **** | a | d |
| Empty *vs. E. coli E48A* | <0.0001 | **** | a | e |
| *E. coli vs. E. teleogrylli* | 0.6940 | ns | b | ab |
| *E. coli vs. R. inusitata* | 0.9970 | ns | b | ab |
| *E. coli vs. R. sp. AA* | 0.4285 | ns | b | ab |
| *E. coli vs. A. sp. B3213* | 0.7681 | ns | b | ab |
| *E. coli vs. P. sp. T1lg23B* | 0.9756 | ns | b | ab |
| *E. coli vs. I. baltica* | 0.0002 | *** | b | c |
| *E. coli vs. E. billingiae* | 0.3931 | ns | b | ab |
| *E. coli vs. P. atlantica* | 0.0017 | ** | b | c |
| *E. coli vs. P. indica* | <0.0001 | **** | b | d |
| *E. coli vs. E. coli E48A* | <0.0001 | **** | b | e |
| *E. teleogrylli vs. R. inusitata* | 0.9955 | ns | ab | ab |
| *E. teleogrylli vs. R. sp. AA* | >0.9999 | ns | ab | ab |
| *E. teleogrylli vs. A. sp. B3213* | >0.9999 | ns | ab | ab |
| *E. teleogrylli vs. P. sp. T1lg23B* | 0.9998 | ns | ab | ab |
| *E. teleogrylli vs. I. baltica* | <0.0001 | **** | ab | c |
| *E. teleogrylli vs. E. billingiae* | >0.9999 | ns | ab | ab |
| *E. teleogrylli vs. P. atlantica* | <0.0001 | **** | ab | c |
| *E. teleogrylli vs. P. indica* | <0.0001 | **** | ab | d |
| *E. teleogrylli vs. E. coli E48A* | <0.0001 | **** | ab | e |
| *R. inusitata vs. R. sp. AA* | 0.9404 | ns | ab | ab |
| *R. inusitata vs. A. sp. B3213* | 0.9986 | ns | ab | ab |
| *R. inusitata vs. P. sp. T1lg23B* | >0.9999 | ns | ab | ab |
| *R. inusitata vs. I. baltica* | <0.0001 | **** | ab | c |
| *R. inusitata vs. E. billingiae* | 0.9229 | ns | ab | ab |
| *R. inusitata vs. P. atlantica* | 0.0002 | *** | ab | c |
| *R. inusitata vs. P. indica* | <0.0001 | **** | ab | d |
| *R. inusitata vs. E. coli E48A* | <0.0001 | **** | ab | e |
| *R. sp. AA vs. A. sp. B3213* | >0.9999 | ns | ab | ab |
| *R. sp. AA vs. P. sp. T1lg23B* | 0.9880 | ns | ab | ab |
| *R. sp. AA vs. I. baltica* | <0.0001 | **** | ab | c |
| *R. sp. AA vs. E. billingiae* | >0.9999 | ns | ab | ab |
| *R. sp. AA vs. P. atlantica* | <0.0001 | **** | ab | c |
| *R. sp. AA vs. P. indica* | <0.0001 | **** | ab | d |
| *R. sp. AA vs. E. coli E48A* | <0.0001 | **** | ab | e |
| *A. sp. B3213 vs. P. sp. T1lg23B* | >0.9999 | ns | ab | ab |
| *A. sp. B3213 vs. I. baltica* | <0.0001 | **** | ab | c |
| *A. sp. B3213 vs. E. billingiae* | >0.9999 | ns | ab | ab |
| *A. sp. B3213 vs. P. atlantica* | <0.0001 | **** | ab | c |
| *A. sp. B3213 vs. P. indica* | <0.0001 | **** | ab | d |

| | | | | |
|---|---|---|---|---|
| *A. sp. B3213 vs. E. coli E48A* | <0.0001 | **** | ab | e |
| *P. sp. T1lg23B vs. I. baltica* | <0.0001 | **** | ab | c |
| *P. sp. T1lg23B vs. E. billingiae* | 0.9822 | ns | ab | ab |
| *P. sp. T1lg23B vs. P. atlantica* | <0.0001 | **** | ab | c |
| *P. sp. T1lg23B vs. P. indica* | <0.0001 | **** | ab | d |
| *P. sp. T1lg23B vs. E. coli E48A* | <0.0001 | **** | ab | e |
| *I. baltica vs. E. billingiae* | <0.0001 | **** | c | ab |
| *I. baltica vs. P. atlantica* | 0.9968 | ns | c | c |
| *I. baltica vs. P. indica* | <0.0001 | **** | c | d |
| *I. baltica vs. E. coli E48A* | <0.0001 | **** | c | e |
| *E. billingiae vs. P. atlantica* | <0.0001 | **** | ab | c |
| *E. billingiae vs. P. indica* | <0.0001 | **** | ab | d |
| *E. billingiae vs. E. coli E48A* | <0.0001 | **** | ab | e |
| *P. atlantica vs. P. indica* | <0.0001 | **** | c | d |
| *P. atlantica vs. E. coli E48A* | <0.0001 | **** | c | e |
| *P. indica vs. E. coli E48A* | 0.0203 | * | d | e |

**Appendix Table S5** p-values from one-way ANOVA Multiple Comparison of Figure 5D

| Comparison | Adjusted p-value | Summary | Letter group 1 | Letter group 2 |
|---|---|---|---|---|
| Empty *vs. E. coli* | 0.0026 | ** | a | b |
| Empty *vs. E. teleogrylli* | 0.2551 | ns | a | ab |
| Empty *vs. R. inusitata* | 0.0239 | * | a | bc |
| Empty *vs. R. sp. AA* | 0.9998 | ns | a | ac |
| Empty *vs. A. sp. B3213* | 0.9989 | ns | a | ac |
| Empty *vs. P. sp. T1lg23B* | 0.0030 | ** | a | b |
| Empty *vs. I. baltica* | <0.0001 | **** | a | b |
| Empty *vs. E. billingiae* | >0.9999 | ns | a | a |
| Empty *vs. P. atlantica* | <0.0001 | **** | a | d |
| Empty *vs. P. indica* | <0.0001 | **** | a | e |
| Empty *vs. E. coli E48A* | <0.0001 | **** | a | f |
| *E. coli vs. E. teleogrylli* | 0.6181 | ns | b | ab |
| *E. coli vs. R. inusitata* | 0.9978 | ns | b | bc |
| *E. coli vs. R. sp. AA* | 0.0148 | * | b | ac |
| *E. coli vs. A. sp. B3213* | 0.0204 | * | b | ac |
| *E. coli vs. P. sp. T1lg23B* | >0.9999 | ns | b | b |
| *E. coli vs. I. baltica* | 0.9587 | ns | b | b |
| *E. coli vs. E. billingiae* | 0.0053 | ** | b | a |
| *E. coli vs. P. atlantica* | 0.0007 | *** | b | d |
| *E. coli vs. P. indica* | <0.0001 | **** | b | e |
| *E. coli vs. E. coli E48A* | <0.0001 | **** | b | f |
| *E. teleogrylli vs. R. inusitata* | 0.9862 | ns | ab | bc |
| *E. teleogrylli vs. R. sp. AA* | 0.6689 | ns | ab | ac |
| *E. teleogrylli vs. A. sp. B3213* | 0.7511 | ns | ab | ac |
| *E. teleogrylli vs. P. sp. T1lg23B* | 0.6559 | ns | ab | b |
| *E. teleogrylli vs. I. baltica* | 0.0646 | ns | ab | b |
| *E. teleogrylli vs. E. billingiae* | 0.4028 | ns | ab | a |
| *E. teleogrylli vs. P. atlantica* | <0.0001 | **** | ab | d |
| *E. teleogrylli vs. P. indica* | <0.0001 | **** | ab | e |
| *E. teleogrylli vs. E. coli E48A* | <0.0001 | **** | ab | f |
| *R. inusitata vs. R. sp. AA* | 0.1136 | ns | bc | ac |
| *R. inusitata vs. A. sp. B3213* | 0.1483 | ns | bc | ac |
| *R. inusitata vs. P. sp. T1lg23B* | 0.9987 | ns | bc | b |
| *R. inusitata vs. I. baltica* | 0.4959 | ns | bc | b |
| *R. inusitata vs. E. billingiae* | 0.0461 | * | bc | a |
| *R. inusitata vs. P. atlantica* | <0.0001 | **** | bc | d |
| *R. inusitata vs. P. indica* | <0.0001 | **** | bc | e |
| *R. inusitata vs. E. coli E48A* | <0.0001 | **** | bc | f |
| *R. sp. AA vs. A. sp. B3213* | >0.9999 | ns | ac | ac |
| *R. sp. AA vs. P. sp. T1lg23B* | 0.0170 | * | ac | b |
| *R. sp. AA vs. I. baltica* | 0.0006 | *** | ac | b |
| *R. sp. AA vs. E. billingiae* | >0.9999 | ns | ac | a |
| *R. sp. AA vs. P. atlantica* | <0.0001 | **** | ac | d |
| *R. sp. AA vs. P. indica* | <0.0001 | **** | ac | e |
| *R. sp. AA vs. E. coli E48A* | <0.0001 | **** | ac | f |
| *A. sp. B3213 vs. P. sp. T1lg23B* | 0.0234 | * | ac | b |
| *A. sp. B3213 vs. I. baltica* | 0.0008 | *** | ac | b |
| *A. sp. B3213 vs. E. billingiae* | >0.9999 | ns | ac | a |
| *A. sp. B3213 vs. P. atlantica* | <0.0001 | **** | ac | d |
| *A. sp. B3213 vs. P. indica* | <0.0001 | **** | ac | e |

| | | | | |
|---|---|---|---|---|
| *A. sp. B3213 vs. E. coli E48A* | <0.0001 | **** | ac | f |
| *P. sp. T1lg23B vs. I. baltica* | 0.9452 | ns | b | b |
| *P. sp. T1lg23B vs. E. billingiae* | 0.0062 | ** | b | a |
| *P. sp. T1lg23B vs. P. atlantica* | 0.0006 | *** | b | d |
| *P. sp. T1lg23B vs. P. indica* | <0.0001 | **** | b | e |
| *P. sp. T1lg23B vs. E. coli E48A* | <0.0001 | **** | b | f |
| *I. baltica vs. E. billingiae* | 0.0002 | *** | b | a |
| *I. baltica vs. P. atlantica* | 0.0187 | * | b | d |
| *I. baltica vs. P. indica* | <0.0001 | **** | b | e |
| *I. baltica vs. E. coli E48A* | <0.0001 | **** | b | f |
| *E. billingiae vs. P. atlantica* | <0.0001 | **** | a | d |
| *E. billingiae vs. P. indica* | <0.0001 | **** | a | e |
| *E. billingiae vs. E. coli E48A* | <0.0001 | **** | a | f |
| *P. atlantica vs. P. indica* | 0.0233 | * | d | e |
| *P. atlantica vs. E. coli E48A* | <0.0001 | **** | d | f |
| *P. indica vs. E. coli E48A* | <0.0001 | **** | e | f |

**Appendix Table S6** p-values from one-way ANOVA Multiple Comparison of Figure 5E

| Comparison | Adjusted p-value | Summary | Letter group 1 | Letter group 2 |
|---|---|---|---|---|
| Empty *vs. E. coli* | 0.7338 | ns | a | a |
| Empty *vs. E. teleogrylli* | >0.9999 | ns | a | a |
| Empty *vs. R. inusitata* | 0.9297 | ns | a | a |
| Empty *vs. R. sp. AA* | >0.9999 | ns | a | a |
| Empty *vs. A. sp. B3213* | >0.9999 | ns | a | a |
| Empty *vs. P. sp. T1lg23B* | 0.0911 | ns | a | ab |
| Empty *vs. I. baltica* | <0.0001 | **** | a | b |
| Empty *vs. E. billingiae* | >0.9999 | ns | a | a |
| Empty *vs. P. atlantica* | 0.0002 | *** | a | b |
| Empty *vs. P. indica* | <0.0001 | **** | a | c |
| Empty *vs. E. coli E48A* | <0.0001 | **** | a | c |
| *E. coli vs. E. teleogrylli* | 0.9614 | ns | a | a |
| *E. coli vs. R. inusitata* | >0.9999 | ns | a | a |
| *E. coli vs. R. sp. AA* | 0.9275 | ns | a | a |
| *E. coli vs. A. sp. B3213* | 0.9595 | ns | a | a |
| *E. coli vs. P. sp. T1lg23B* | 0.9538 | ns | a | ab |
| *E. coli vs. I. baltica* | 0.0038 | ** | a | b |
| *E. coli vs. E. billingiae* | 0.8255 | ns | a | a |
| *E. coli vs. P. atlantica* | 0.0184 | * | a | b |
| *E. coli vs. P. indica* | <0.0001 | **** | a | c |
| *E. coli vs. E. coli E48A* | <0.0001 | **** | a | c |
| *E. teleogrylli vs. R. inusitata* | 0.9981 | ns | a | a |
| *E. teleogrylli vs. R. sp. AA* | >0.9999 | ns | a | a |
| *E. teleogrylli vs. A. sp. B3213* | >0.9999 | ns | a | a |
| *E. teleogrylli vs. P. sp. T1lg23B* | 0.2617 | ns | a | ab |
| *E. teleogrylli vs. I. baltica* | 0.0001 | *** | a | b |
| *E. teleogrylli vs. E. billingiae* | >0.9999 | ns | a | a |
| *E. teleogrylli vs. P. atlantica* | 0.0007 | *** | a | b |
| *E. teleogrylli vs. P. indica* | <0.0001 | **** | a | c |
| *E. teleogrylli vs. E. coli E48A* | <0.0001 | **** | a | c |
| *R. inusitata vs. R. sp. AA* | 0.9938 | ns | a | a |
| *R. inusitata vs. A. sp. B3213* | 0.9979 | ns | a | a |
| *R. inusitata vs. P. sp. T1lg23B* | 0.7873 | ns | a | ab |
| *R. inusitata vs. I. baltica* | 0.0013 | ** | a | b |
| *R. inusitata vs. E. billingiae* | 0.9681 | ns | a | a |
| *R. inusitata vs. P. atlantica* | 0.0067 | ** | a | b |
| *R. inusitata vs. P. indica* | <0.0001 | **** | a | c |
| *R. inusitata vs. E. coli E48A* | <0.0001 | **** | a | c |
| *R. sp. AA vs. A. sp. B3213* | >0.9999 | ns | a | a |
| *R. sp. AA vs. P. sp. T1lg23B* | 0.2059 | ns | a | ab |
| *R. sp. AA vs. I. baltica* | 0.0001 | *** | a | b |
| *R. sp. AA vs. E. billingiae* | >0.9999 | ns | a | a |
| *R. sp. AA vs. P. atlantica* | 0.0005 | *** | a | b |
| *R. sp. AA vs. P. indica* | <0.0001 | **** | a | c |
| *R. sp. AA vs. E. coli E48A* | <0.0001 | **** | a | c |
| *A. sp. B3213 vs. P. sp. T1lg23B* | 0.2574 | ns | a | ab |
| *A. sp. B3213 vs. I. baltica* | 0.0001 | *** | a | b |
| *A. sp. B3213 vs. E. billingiae* | >0.9999 | ns | a | a |
| *A. sp. B3213 vs. P. atlantica* | 0.0007 | *** | a | b |
| *A. sp. B3213 vs. P. indica* | <0.0001 | **** | a | c |

| | | | | |
|---|---|---|---|---|
| *A. sp. B3213 vs. E. coli E48A* | <0.0001 | **** | a | c |
| *P. sp. T1lg23B vs. I. baltica* | 0.0879 | ns | ab | b |
| *P. sp. T1lg23B vs. E. billingiae* | 0.1276 | ns | ab | a |
| *P. sp. T1lg23B vs. P. atlantica* | 0.3024 | ns | ab | b |
| *P. sp. T1lg23B vs. P. indica* | <0.0001 | **** | ab | c |
| *P. sp. T1lg23B vs. E. coli E48A* | <0.0001 | **** | ab | c |
| *I. baltica vs. E. billingiae* | <0.0001 | **** | b | a |
| *I. baltica vs. P. atlantica* | 0.9999 | ns | b | b |
| *I. baltica vs. P. indica* | <0.0001 | **** | b | c |
| *I. baltica vs. E. coli E48A* | <0.0001 | **** | b | c |
| *E. billingiae vs. P. atlantica* | 0.0003 | *** | a | b |
| *E. billingiae vs. P. indica* | <0.0001 | **** | a | c |
| *E. billingiae vs. E. coli E48A* | <0.0001 | **** | a | c |
| *P. atlantica vs. P. indica* | <0.0001 | **** | b | c |
| *P. atlantica vs. E. coli E48A* | <0.0001 | **** | b | c |
| *P. indica vs. E. coli E48A* | 0.3207 | ns | c | c |

**Appendix Table S7** p-values from one-way ANOVA Multiple Comparison of Figure 5F

| Comparison | Adjusted p-value | Summary | Letter group 1 | Letter group 2 |
|---|---|---|---|---|
| Empty *vs. E. coli* | 0.1809 | ns | a | a |
| Empty *vs. E. teleogrylli* | 0.9998 | ns | a | a |
| Empty *vs. R. inusitata* | 0.4125 | ns | a | a |
| Empty *vs. R. sp. AA* | >0.9999 | ns | a | a |
| Empty *vs. A. sp. B3213* | 0.8074 | ns | a | a |
| Empty *vs. P. sp. T1lg23B* | 0.9984 | ns | a | a |
| Empty *vs. I. baltica* | <0.0001 | **** | a | b |
| Empty *vs. E. billingiae* | >0.9999 | ns | a | a |
| Empty *vs. P. atlantica* | <0.0001 | **** | a | b |
| Empty *vs. P. indica* | <0.0001 | **** | a | c |
| Empty *vs. E. coli E48A* | <0.0001 | **** | a | d |
| *E. coli vs. E. teleogrylli* | 0.5258 | ns | a | a |
| *E. coli vs. R. inusitata* | >0.9999 | ns | a | a |
| *E. coli vs. R. sp. AA* | 0.2832 | ns | a | a |
| *E. coli vs. A. sp. B3213* | 0.9865 | ns | a | a |
| *E. coli vs. P. sp. T1lg23B* | 0.6531 | ns | a | a |
| *E. coli vs. I. baltica* | 0.0010 | *** | a | b |
| *E. coli vs. E. billingiae* | 0.1996 | ns | a | a |
| *E. coli vs. P. atlantica* | 0.0073 | ** | a | b |
| *E. coli vs. P. indica* | <0.0001 | **** | a | c |
| *E. coli vs. E. coli E48A* | <0.0001 | **** | a | d |
| *E. teleogrylli vs. R. inusitata* | 0.8281 | ns | a | a |
| *E. teleogrylli vs. R. sp. AA* | >0.9999 | ns | a | a |
| *E. teleogrylli vs. A. sp. B3213* | 0.9923 | ns | a | a |
| *E. teleogrylli vs. P. sp. T1lg23B* | >0.9999 | ns | a | a |
| *E. teleogrylli vs. I. baltica* | <0.0001 | **** | a | b |
| *E. teleogrylli vs. E. billingiae* | >0.9999 | ns | a | a |
| *E. teleogrylli vs. P. atlantica* | <0.0001 | **** | a | b |
| *E. teleogrylli vs. P. indica* | <0.0001 | **** | a | c |
| *E. teleogrylli vs. E. coli E48A* | <0.0001 | **** | a | d |
| *R. inusitata vs. R. sp. AA* | 0.5710 | ns | a | a |
| *R. inusitata vs. A. sp. B3213* | >0.9999 | ns | a | a |
| *R. inusitata vs. P. sp. T1lg23B* | 0.9117 | ns | a | a |
| *R. inusitata vs. I. baltica* | 0.0003 | *** | a | b |
| *R. inusitata vs. E. billingiae* | 0.4443 | ns | a | a |
| *R. inusitata vs. P. atlantica* | 0.0021 | ** | a | b |
| *R. inusitata vs. P. indica* | <0.0001 | **** | a | c |
| *R. inusitata vs. E. coli E48A* | <0.0001 | **** | a | d |
| *R. sp. AA vs. A. sp. B3213* | 0.9172 | ns | a | a |
| *R. sp. AA vs. P. sp. T1lg23B* | >0.9999 | ns | a | a |
| *R. sp. AA vs. I. baltica* | <0.0001 | **** | a | b |
| *R. sp. AA vs. E. billingiae* | >0.9999 | ns | a | a |
| *R. sp. AA vs. P. atlantica* | <0.0001 | **** | a | b |
| *R. sp. AA vs. P. indica* | <0.0001 | **** | a | c |
| *R. sp. AA vs. E. coli E48A* | <0.0001 | **** | a | d |
| *A. sp. B3213 vs. P. sp. T1lg23B* | 0.9986 | ns | a | a |
| *A. sp. B3213 vs. I. baltica* | <0.0001 | **** | a | b |
| *A. sp. B3213 vs. E. billingiae* | 0.8346 | ns | a | a |
| *A. sp. B3213 vs. P. atlantica* | 0.0004 | *** | a | b |
| *A. sp. B3213 vs. P. indica* | <0.0001 | **** | a | c |

| | | | | |
|---|---|---|---|---|
| *A. sp. B3213 vs. E. coli E48A* | <0.0001 | **** | a | d |
| *P. sp. T1lg23B vs. I. baltica* | <0.0001 | **** | a | b |
| *P. sp. T1lg23B vs. E. billingiae* | 0.9991 | ns | a | a |
| *P. sp. T1lg23B vs. P. atlantica* | <0.0001 | **** | a | b |
| *P. sp. T1lg23B vs. P. indica* | <0.0001 | **** | a | c |
| *P. sp. T1lg23B vs. E. coli E48A* | <0.0001 | **** | a | d |
| *I. baltica vs. E. billingiae* | <0.0001 | **** | b | a |
| *I. baltica vs. P. atlantica* | 0.9992 | ns | b | b |
| *I. baltica vs. P. indica* | 0.0003 | *** | b | c |
| *I. baltica vs. E. coli E48A* | <0.0001 | **** | b | d |
| *E. billingiae vs. P. atlantica* | <0.0001 | **** | a | b |
| *E. billingiae vs. P. indica* | <0.0001 | **** | a | c |
| *E. billingiae vs. E. coli E48A* | <0.0001 | **** | a | d |
| *P. atlantica vs. P. indica* | <0.0001 | **** | b | c |
| *P. atlantica vs. E. coli E48A* | <0.0001 | **** | b | d |
| *P. indica vs. E. coli E48A* | <0.0001 | **** | c | d |

**Appendix Table S8** Uniprot RfaH genomic context

| Clade | Clade features | Number of found genomic contexts | Next to operon | LPS in operon |
|---|---|---|---|---|
| 1 | - | 0 | - | - |
| 2 | Halophilic gammaproteobacteria | 5 | 3 | 0 |
| 4 | Pseudomonadaceae | 8 | 3 | 1 |
| 5 | Lithotrophic gammaproteobacteria | 6 | 2 | 0 |
